# Model-driven experimental design identifies counter-acting feedback regulation in the osmotic stress response of yeast

**DOI:** 10.1101/2020.04.20.051599

**Authors:** Sara Kimiko Suzuki, Beverly Errede, Henrik G. Dohlman, Timothy C. Elston

## Abstract

Cells rely on mitogen-activated protein kinases (MAPKs) to survive environmental stress. In yeast, activation of the MAPK Hog1 is known to mediate the response to high osmotic conditions. Recent studies of Hog1 revealed that its temporal activity is subject to both negative and positive feedback regulation, yet the mechanisms of feedback remain unclear. By designing mathematical models of increasing complexity for the Hog1 MAPK cascade, we identified pathway circuitry sufficient to capture Hog1 dynamics observed in vivo. We used these models to optimize experimental designs for distinguishing potential feedback loops. Performing experiments based on these models revealed mutual inhibition between Hog1 and its phosphatases as the likely positive feedback mechanism underlying switch-like, dose-dependent MAPK activation. Importantly, our findings reveal a new signaling function for MAPK phosphatases. More broadly, they demonstrate the value using mathematical models to infer targets of feedback regulation in signaling pathways.

## Introduction

All cells rely on intracellular signaling systems to protect themselves from environmental stress. These pathways execute the appropriate cellular response by relaying the strength, duration, and other quantitative information about changing environmental conditions (Alon, 2007; Purvis and Lahav, 2013). When the external stimulus is harmful to the cell, the cell’s response can determine whether it survives. To mitigate the effects of stress, cells use signaling pathways that often incorporate mitogen-activated protein kinase (MAPK) cascades (Cargnello and Roux, 2011). In this three-tiered signaling motif, a MAPK kinase kinase (MAP3K) phosphorylates a MAPK kinase (MAP2K), which in turn phosphorylates a terminal MAPK. These phosphorylation events occur within the activation loop of the kinase domain, thereby enabling catalytic activity. The MAPK coordinates all events required for a proper response to the environmental stress.

The MAPK signaling cascade is conserved in all eukaryotic organisms, from humans to yeast. In the case of the *S. cerevisiae* High Osmolarity Glycerol (HOG) pathway, hyperosmotic stress initiates signaling to activate cytoprotective responses (Brewster and Gustin, 2014; Miermont et al., 2011; O’Rourke et al., 2002; Saito and Posas, 2012). This signaling occurs through the Sln1 and Sho1 input branches where the Sln1 branch has two MAP3Ks, Ssk2 and Ssk22, while the Sho1 branch has a single MAP3K, Ste11 (Figure 1). All three of these MAP3Ks converge on and activate the MAP2K Pbs2, while Pbs2 alone activates the MAPK Hog1. Hog1 protects the cell by increasing cytosolic osmolyte concentrations to reestablish turgor pressure over time. Hog1 is known to phosphorylate at least 35 proteins, some of which are transcription factors in the nucleus leading to the induction or repression of ∼300 genes (Capaldi et al., 2008; Janschitz et al., 2019; O’Rourke and Herskowitz, 2004). Activation of this pathway is transient, and once cells have fully-adapted, they can resume cell cycle progression and proliferation (Escoté et al., 2011).

**Figure 1:**
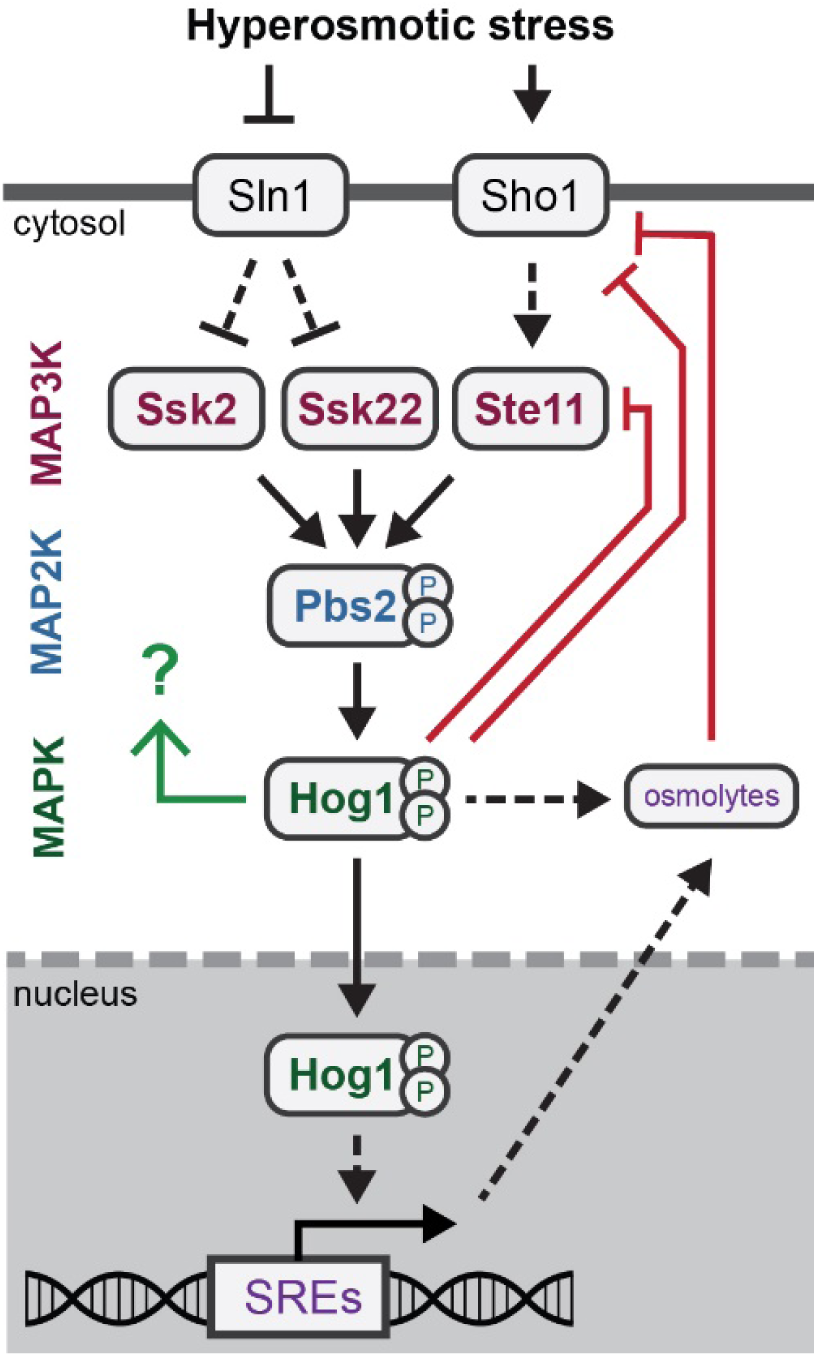
Hog1-dependent feedbacks within the HOG Pathway. Two input branches activate a MAPK cascade to initiate adaptation to hyperosmotic stress. Hog1 controls its own phosphorylation dynamics through negative (red arrows) and positive (green arrow) feedback mechanisms. Hog1 phosphorylates upstream HOG pathway components, including Ste11, Ssk2, and Sho1, which downregulates signaling. Hog1 increases osmolyte concentrations by cytosolic and nuclear events, such as the closing of glycerol export channels and the transcription of genes with Stress Response Elements (SREs). The increase of intracellular osmolarity also suppresses HOG signaling, putatively at the level of receptors. Finally, Hog1 likely initiates a positive feedback loop, but the target is still unknown.

Many MAPK signaling networks rely on feedback regulation to amplify or diminish a signal in a time-dependent manner (Albeck et al., 2013; Brandman and Meyer, 2008). While multiple feedback loops have been identified for the HOG pathway, it is unknown how these regulatory mechanisms function together or if they are sufficient to capture the dynamics of Hog1 activity. For example, Hog1 phosphorylates the osmosensor Sho1 and the MAP3K Ssk2, leading to diminished signal transduction (Hao et al., 2007; Sharifian et al., 2015). Similarly, Hog1-dependent phosphorylation of Ste50, the adapter protein of Ste11, increases Ste50 dissociation from its signaling complex, thereby downregulating signal transmission (Nagiec and Dohlman, 2012; Yamamoto et al., 2010). Hog1-dependent activity also attenuates signaling by initiating the closure of osmolyte channels, inducing the synthesis of amino acid metabolites, increasing the production of osmolytes like glycerol and trehalose, and inducing the transcription of osmolyte metabolism-associated genes (Babazadeh et al., 2014; Lee et al., 2013; O’Rourke and Herskowitz, 2004; Petelenz-Kurdziel et al., 2013; Shellhammer et al., 2017). Phosphorylation and osmolyte accumulation act on different timescales to suppress Hog1 signaling, and therefore could differentially affect the dynamics of Hog1 activity.

Further studies have shown that Hog1 acts, in part, by regulating its own catalytic activity. In response to a wide range of external salt concentrations, Hog1 is rapidly and fully phosphorylated, whereas dephosphorylation occurs at increasingly later times as the dose of the stimulus increases (English et al., 2015). Because different doses of salt elicit different transcriptional responses, it is likely that signal duration, rather than amplitude, transmits information regarding the external salt concentration. This behavior has been referred to as “dose-to-duration” signaling (Behar et al., 2008). Experiments using a Hog1 variant (Hog1-as) engineered to respond to a pharmacological inhibitor (Klein et al., 2011; Westfall and Thorner, 2006) revealed that Hog1 kinase activity affects its response to high osmotic stress. In the absence of MAPK kinase activity, the maximal level of Hog1 phosphorylation is dependent on the concentration of external osmolytes; peak phosphorylation is delayed and is far more sustained than that of the wildtype MAPK (English et al., 2015). These observations indicate that Hog1 kinase activity is required for full activation by the MAP2K Pbs2 and for timely inactivation by appropriate phosphatases. Such behaviors are indicative of positive and negative feedback. However, while the necessity of feedback within the HOG pathway has long been appreciated, many details of the mechanisms controlling Hog1 phosphorylation dynamics are still unknown. This is in part due to the complexities of the observed behaviors, which are both dose- and time-dependent.

One approach to understanding complex biological data is to describe them using mathematical models. The structures of HOG pathway models have varied substantially, from having minimal two state systems to representing all of the HOG pathway components (Klipp and Schaber, 2008; Mitchell et al., 2015; Stojanovski et al., 2017). A subset has focused on negative feedback while others have investigated the role of the two input branches (Granados et al., 2017; Hersen et al., 2008; Schaber et al., 2012). Many models investigating feedback regulation concluded that the pathway needs Hog1-dependent integral negative feedback control to exhibit perfect adaptation (Klipp et al., 2005; Mettetal et al., 2008; Mitchell et al., 2015; Muzzey et al., 2009; Zi et al., 2010). Other models further explored different mechanisms of negative feedback, proposing that the required feedback mechanism entails the slow accumulation of osmolytes (Petelenz-Kurdziel et al., 2013; Schaber et al., 2012). Subsequently, our experimental efforts revealed the potential importance of positive feedback for fast Hog1 activation (English et al., 2015). However, there are no reported mechanisms of positive feedback for the HOG pathway.

Despite substantial progress, a complete systems-level understanding for the role of counter-acting feedback regulation is still lacking. Therefore, our goal here was to perform a systematic computational analysis of Hog1 activity that could identify likely targets of feedback regulation and to then design experiments to test the predicted feedback loops. The starting point for our investigations was a model of a three-tiered MAPK cascade to which we systematically added different potential feedback motifs. In particular, we started with a minimal model for adaptation involving a single negative feedback loop (Ferrell, 2016), and added candidate feedback loops until we were able to reproduce Hog1 phosphorylation dynamics. Combining both modeling and biological experiments allowed us to identify the necessary feedback mechanisms by using each method to inform the other in an iterative process. As detailed below, our investigations determined that fast positive feedback and delayed negative feedback can account for the time- and dose-dependent behaviors of Hog1. Further analysis suggests that positive feedback is controlled by Hog1 down regulation of its phosphatase activity.

## Results

### Hog1 and Pbs2 phosphorylation are dependent on Hog1 kinase activity

Our broad objective is to identify pathway circuitry for regulating MAPK signaling generally, and for the HOG pathway in particular. We first collected the experimental data depicting pathway dynamics. Our approach was to design time-course experiments that measure the dynamics of Hog1 and its upstream kinase, Pbs2, under various experimental conditions. Using this approach, we defined 10 important characteristics of the HOG pathway that our models need to capture in order to be biologically accurate. These 10 characteristics are enumerated in the following section and are summarized below.

We first assessed Hog1 phosphorylation upon hyperosmotic stress. We exposed liquid cultures to a range of KCl concentrations and collected whole-cell lysates over time. To quantify the proportion of phosphorylated Hog1, we used Phos-tag immunoblotting, which resolves different states of a protein in proportion to the number of sites phosphorylated (Kinoshita et al., 2006). Because wildtype Hog1 is normally either unphosphorylated or dually phosphorylated, we can easily distinguish the two forms of the protein after SDS-PAGE with the Phos-tag reagent (Figure 2A, left). The stoichiometry of phosphorylation was calculated as the proportion of dually phosphorylated Hog1 compared to the total amount of Hog1 in each lane (Figure 2A, right). Consistent with previous results (English et al., 2015), we observed three characteristic features of Hog1: (1) no basal activation, (2) fast and full activation in response to KCl, and (3) transient duration of activation that is proportional to the KCl dose. These features give rise to dose-to-duration signaling.

**Figure 2:**
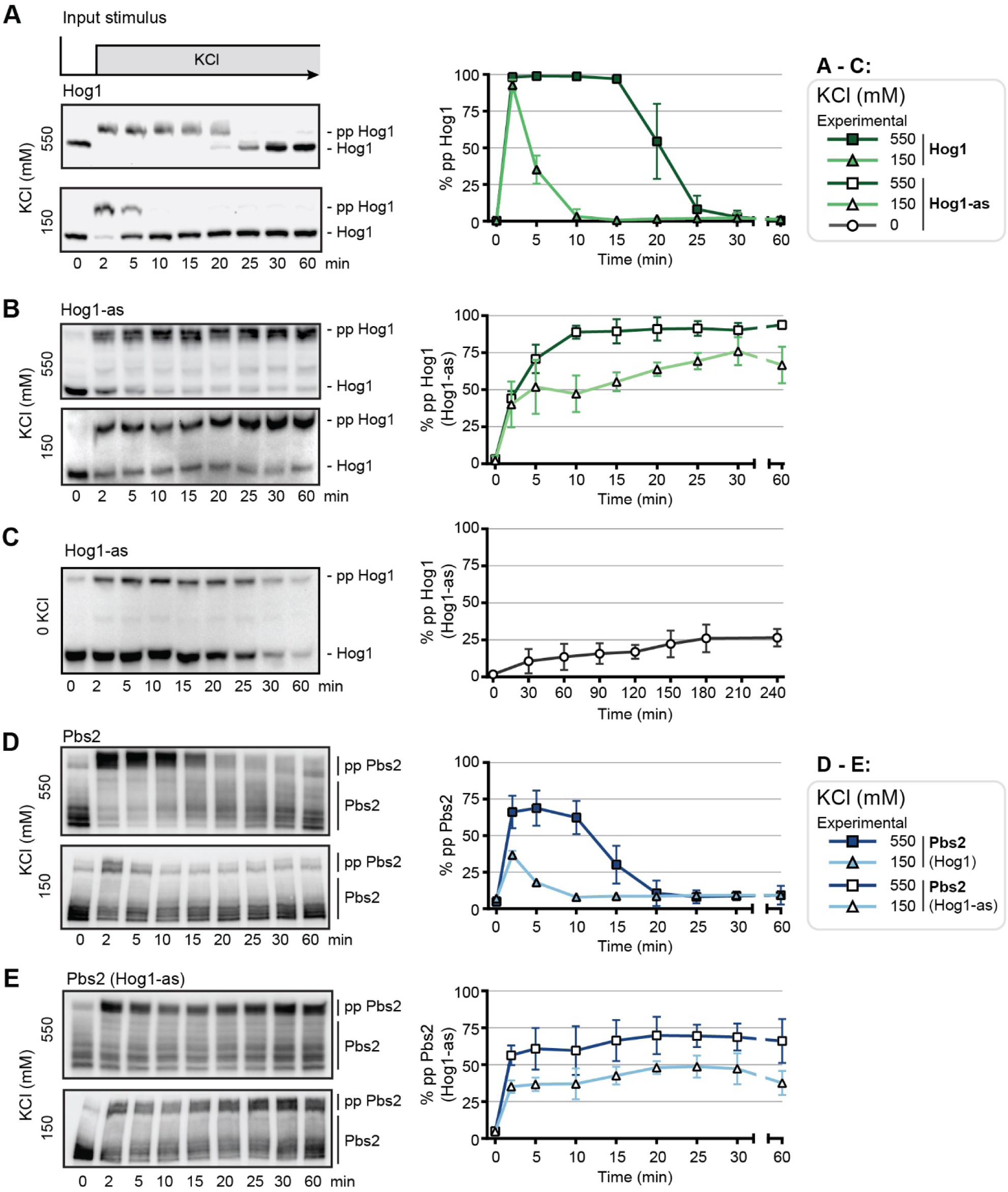
HOG pathway dynamics. **(A)** Left: Hog1 dual phosphorylation (pp Hog1) over time in response to a single step stimulus (top) of 550 mM KCl (center) or 150 mM KCl (bottom), resolved using the Phos-tag method. Right: Quantification of blots. **(B)** Same as (A) but using an analog sensitive Hog1 + ATP analog (Hog1-as). **(C)** Same as (B) but taken for longer time points and in the absence of KCl. **(D)** Left: Pbs2 phosphorylation over time in response to 550 mM KCl (top) and 150 mM KCl (bottom), resolved using the Phos-tag method. **(E)** Same as (D) but using Hog1-as. Error bars represent SD of each point. All experimental data are n = 3.

Hog1 kinase activity can be selectively blocked using an analog-sensitive Hog1 variant (Hog1 ^T100A^) that is inhibited with the ATP analog, [1-(1,1-dimethylethyl)-3-(1-naphthalenyl)1H-pyrazolo[3,4-d]pyrimidin-4-amine] or 1-NA-PP1 (Kung et al., 2006). Accordingly, we stimulated cells following Hog1^T100A^ inhibition, and ran Phos-tag SDS-PAGE as previously described (English et al., 2015). Inhibited Hog1 (Hog1-as) exhibited three characteristics that differ from wildtype. Consistent with results of English et al. (2015), we observed Hog1 dynamics that were: (4) slow and (5) sustained (Figure 2B). The slowed rate of activation and lack of full dual phosphorylation when Hog1 is inhibited indicates the presence of Hog1-dependent positive feedback. Furthermore, as noted previously, Hog1 exhibits basal dual phosphorylation when its kinase activity is inhibited (English et al., 2015; Macia et al., 2009; Schaber et al., 2012). Our quantification revealed that under these conditions dually phosphorylated Hog1 slowly accumulates, reaching a steady state of approximately 30% of the total (characteristic feature (6)) (Figure 2C). The lack of signal attenuation and increase in the basal level of dually phosphorylated Hog1 in the absence of Hog1 activity demonstrate Hog1-dependent negative feedback. These results reflect the complexity of HOG signaling and motivate our investigations to determine where in the pathway feedback regulation acts.

We next measured the dynamics of another upstream signaling component, at multiple salt concentrations, with and without Hog1 kinase inhibition. Our rationale was that these experiments would provide important additional data for informing our models and identifying targets of feedback regulation. We chose the MAP2K Pbs2 because it is more abundant than any one of the MAP3Ks, is common to both input branches of the pathway and is phosphorylated when activated (Tatebayashi et al., 2020). Thus, we used the Phos-tag western blotting technique described above to measure the dose- and Hog1-kinase dependency of Pbs2 phosphorylation dynamics. As shown in Figure 2D, osmotic stress stimulation caused a mobility shift of Pbs2 that was (7) fast and partial and also (8) transient. This behavior mirrored two of the wildtype properties, but unlike Hog1, Pbs2 did not become fully phosphorylated at 150 mM KCl. When Hog1 was kinase-inhibited, Pbs2 phosphorylation was also (9) fast and partial, but (10) sustained, as observed for Hog1-as, indicating signal attenuation occurs earlier in the pathway.

### Delayed negative feedback promotes pathway deactivation

Our next step was to identify potential HOG feedback circuits by fitting models to our Hog1 and Pbs2 phosphorylation data. We considered a model successful if it could capture the 10 pathway characteristics enumerated above. We took a systematic approach by beginning with a minimal model for adaptive behavior and adding complexity as needed. In this way, we hoped to gain insight to the limitations of each model. The minimal model (Model I) for the HOG MAPK cascade is comprised of a single negative feedback loop initiating from Hog1 and targeting the MAP3K (Figure 3A). From a biological perspective, this model represents Hog1 suppressing its own activity by diminishing the rate at which the MAPK3K is activated, this might occur through increasing the intracellular osmolyte concentration, feedback phosphorylation or both (English et al., 2015; Hao et al., 2007; Sharifian et al., 2015). The model consists of three species, representing each of the three kinases in the MAPK cascade. We modeled phosphorylation and dephosphorylation using Michaelis-Menten kinetics and ignored the synthesis and degradation of the kinases, as their expression is not known to be induced following hyperosmotic stress.

**Figure 3:**
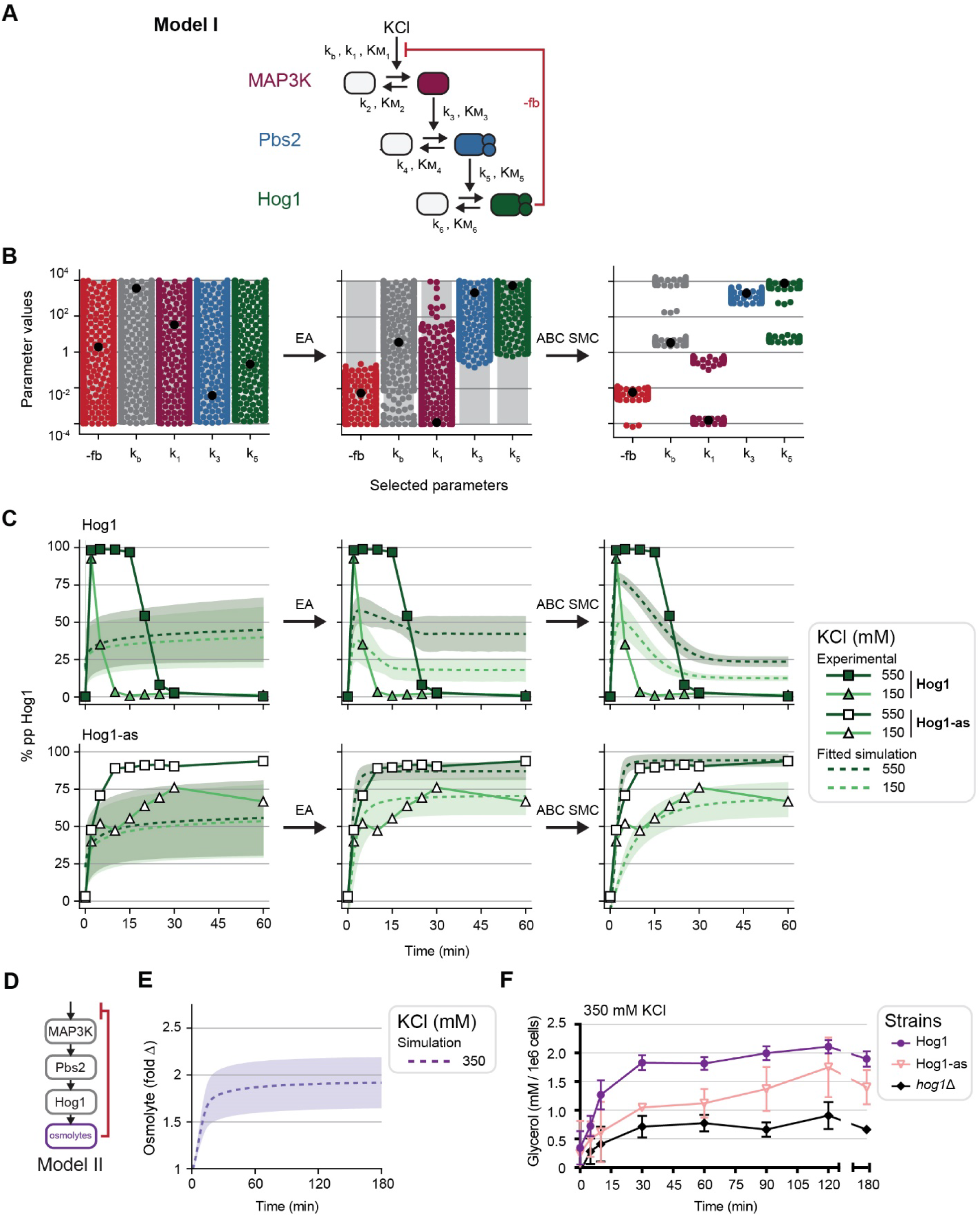
Model building and parameter estimation of potential feedback circuits. **(A)** Schematic of Model I, a single negative feedback from Hog1, targeting the input with associated parameters to be estimated. **(B)** The parameter optimization method. First, parameter values are randomly assigned, then the Evolutionary Algorithm (EA) finds candidate parameter sets, and finally, the Approximate Bayesian Computation Sequential Monte Carlo (ABC SMC) searches the local parameter space surrounding the EA parameter sets. Gray bars indicate the range of potential values selected uniformly during the EA. Colored points specify parameter values and black points highlight the best (lowest MSE between experimental data and simulations) parameter values after each step. **(C)** Simulated fits at each estimation step are overlaid with wildtype Hog1 (top row, filled symbols) and Hog1-as (bottom row, open symbols) data at each parameter optimization step. Average simulated behaviors are plotted using dashed lines. All simulations are n = 1000 and all shaded regions are a SD of 1. **(D)** Schematic of Model II that features a delayed negative feedback, presumably from osmolyte accumulation. **(E)** Model II simulated prediction of downstream component behavior. **(F)** Glycerol accumulation over time in response to 350 mM KCl with and without Hog1 activity. *hog1*Δ cells served as a negative control. All experiments are n = 3 and error bars represent SD of each point.

*Model I* (Figure 3A):

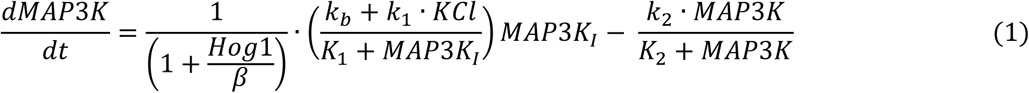

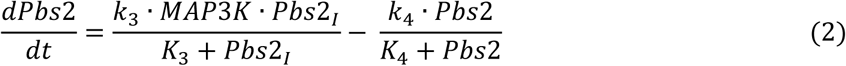

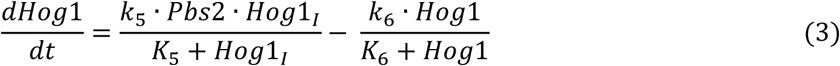

where each K_i_ represents a Michaelis constant, the k_i_’s are either the k_cat_ or V_max_ of the reaction, depending on whether the enzyme concentration is explicitly taken into account, and k_b_ is the basal activation rate (English et al., 2015; Macia et al., 2009). We assumed that salt increases the V_max_ of the reaction for activation of the MAP3K. That is, k_1_ = k_1_’ KCl. We used a decreasing Hill function to include negative feedback, with β representing the concentration of active Hog1 needed to reduce the MAP3K activation rate by half.

Having defined a model, we then sought to fit it to our experimental data. To perform parameter estimation, we created a hybrid method, combining a global and local search method that minimized the distance between simulated fits and the data based on mean squared error (MSE). Recent benchmark efforts have shown that similar combination strategies are particularly efficient and best performing when compared to stand alone methods (Villaverde et al., 2019). Here, we used an evolutionary algorithm (EA) (Fortin et al., 2012) that performs a global search method within a large search space to find best-fitting parameter sets. We then used an approximate Bayesian computation sequential Monte Carlo (ABC SMC) method (Toni et al., 2009) to fine-tune the EA-determined parameter sets to further realize distributions of model parameter values that produced results consistent with the data (Figure 3B-C). This process resulted in 1000 parameter sets that could meet the model-specific MSE thresholds. Details of our parameter optimization method are provided in the Methods section.

We fit Model I to our data presented in Figure 2. We simulated wildtype behavior with the full system and simulated kinase-inhibited behavior by removing the Hog1-dependent negative feedback loop. As shown in Figure 3C (top right panel), Model I could neither capture full Hog1 activation nor full deactivation. We inferred that Model I was unsuccessful at capturing our data because in the model Hog1 activity immediately suppresses activation of the MAP3K, and consequentially the model cannot simultaneously satisfy the constraints of full activation in wildtype cells and the amplitude dependence of Hog1-as strain. This reasoning is in agreement with work in Schaber et al. 2012 in which they found a delayed negative feedback is necessary for full signal attenuation while negative feedback originating from Hog1 serves to fine-tune the response. Hence, we expanded Model I to include an additional step between Hog1 activation and pathway inhibition to produce a time delay in the negative feedback loop (Figure 3D). This circuitry is consistent with prior models (Ma et al., 2009) and experimental studies, which demonstrate that full HOG pathway adaptation requires an increase in cytosolic osmolytes (Babazadeh et al., 2014; Hohmann, 2002; Siderius et al., 2000), though this model species could represent any upregulated signaling processes downstream of Hog1. Therefore, we updated the model to include this process.

*Model II* (Figure 3D):

Model I equations 2, and 3 remain the same in Model II. The osmolyte concentration was modeled using the following equation:

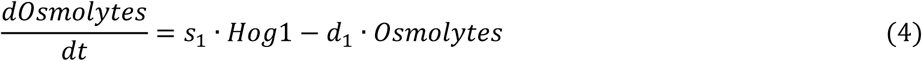

where s_1_ is the rate of osmolyte synthesis, which requires active Hog1, and d_1_ is rate of osmolyte degradation. Model I equation 1 was updated to replace active Hog1 in the negative feedback term with the osmolyte concentration. Overall, adding a delay in the negative feedback loop significantly improved the performance of the model and allowed it to capture full Hog1 deactivation (Figure 3 – figure supplement 1). However, Model II still could not capture full Hog1 phosphorylation.

Investigating the behavior of the osmolyte concentration, we found that Model II predicted a 2-fold increase of the putative osmolyte species over the course of 15 minutes after 350 mM KCl stimulus (Figure 3E). To test the model, we exposed cells, with and without Hog1 activity, with 350 mM KCl and measured glycerol accumulation over time (Figure 3E). While glycerol exhibited a higher-than predicted increase (four-fold vs two-fold), the dynamics were similar to the model prediction (Figure 3F). The discrepancy in abundance is likely due to Hog1-independent glycerol production. We observed 1- to 2-fold increase of glycerol accumulation in the *hog1*Δ and Hog1-as cells treated with 1-NA-PP1 (Figure 3E, and Figure 3 – figure supplement 2), as reported previously for glycerol and other osmolytes (Babazadeh et al., 2014; Petelenz-Kurdziel et al., 2013; Shellhammer et al., 2017). We then perturbed the behavior of Model II’s osmolyte species to understand how the osmolyte concentration controlled Hog1 dynamics. Increasing the osmolyte synthesis rate caused the osmolytes to accumulate faster than the fitted simulations which limited Hog1 activation to 5 minutes (Figure 3 – figure supplement 3, left compared to center). Decreasing the osmolyte synthesis rate caused it to accumulate more slowly thereby extending the duration of Hog1 activation to over an hour (Figure 3 – figure supplement 3, left compared to right). This delayed negative feedback does not only control the timing of Hog1 phosphorylation, but also the ability of Hog1 to fully adapt, as seen when decreasing the osmolyte synthesis rate (Figure 3 – figure supplement 3, bottom right). Altogether, these data suggest that a necessary negative feedback originates from a downstream species for full signal attenuation, and that species could likely be the accumulation of intracellular osmolytes.

Thus, compared to Model I, Model II was better able to capture Hog1 dose-to duration dynamics and Pbs2 dynamics. However, the revised model still failed to capture full Hog1 activation and poorly replicated other features of the data, such as the basal phosphorylation dynamics in the Hog1-as strain. While at this point, we do not rule out Model II from further consideration, its inability to replicate several pathway features motivated us to investigate if other potential feedback loops.

### Fast positive feedback promotes pathway activation

Model II captured many of the characteristics of Hog1 activation and deactivation. However, Model II did not reach full activation of Hog1, even at the highest concentrations of stimulus. This failure of the model suggests that it lacks an important positive feedback loop. Since Hog1 activation occurs within two minutes, positive feedback would need to act rapidly. Thus, we hypothesized that it originates from Hog1 directly phosphorylating a pathway component. To test this possibility, we expanded Model II into three new models (Models IIa-c) that include Hog1-driven positive feedback loops targeting one of the three kinases in the MAPK kinase cascade: ‘a’ targets the MAP3K, ‘b’ targets Pbs2, and ‘c’ targets Hog1 itself. These loops were modeled by including a term in the relevant activation rate that was proportional to the level of active Hog1.

For example, *Model IIc* includes Model II equations 1, 2, 3, and 4 with the following modification to the equation for Hog1:

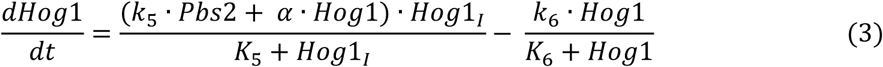

where Hog1-mediated positive feedback (*α*. *Hog*1) increases its own activation. We used the same procedure as described above to train the models. A summary of each model’s fit to the data is provided in Figure 4A. Model IIa produced results very similar to Model II (Figure 4 – figure supplement 1), while Models IIb-c with positive feedback targeting Pbs2 and Hog1, respectively, produced better fits to the data and were able to capture all 10 pathway characteristics (Model IIb: Figure 4 – figure supplement 2, Model IIc: Figure 4B-D). We also found that these two models could predict wildtype Hog1 behavior in response to intermediate single-step KCl concentrations: 250, 350, and 450 mM KCl from in English et al., 2015 (Model IIb: Figure 4 – figure supplement 2, Model IIc: Figure 4E (left)). These models also followed similar Hog1-as dynamics as the previously published though the previously published data is slightly higher than that seen in our data (Figure 4E, right compared to Figure 4D, center). Even with this small discrepancy, these data suggest Hog1 phosphorylates a pathway component at or below that of the MAP2K Pbs2, forming a positive feedback loop.

**Figure 4:**
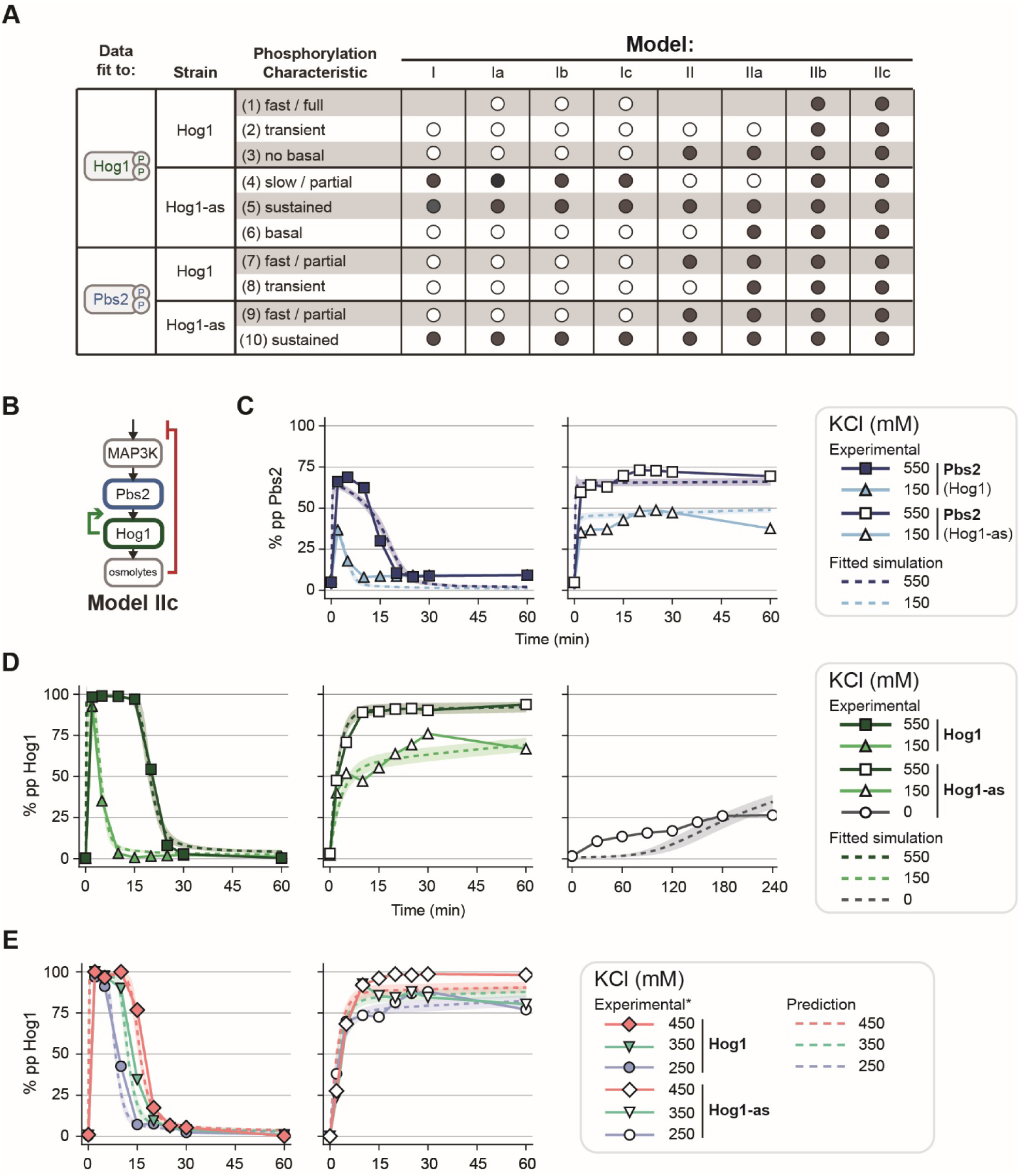
Model fits and predictions to single step stimuli. **(A)** Table showing model fits to each of the HOG pathway characteristics. Dots indicate that the model captures the behavior, where filled circles fit the experimental data well and hollow circles do not. **(B)** Schematic of one of the two models that fits all of the phosphorylation characteristics. **(C)** Model IIc simulated Pbs2 fits (dashed lines) overlaid with experimental data (symbols). Left: Data and simulations for wildtype Hog1 in response to 550 mM and 150 mM KCl. Right: Data and simulations for Hog1-as. **(D)** Model IIc simulated Hog1 fits (dashed lines) overlaid with experimental data (symbols). Left: Data and simulations for wildtype Hog1 in response to 550 mM and 150 mM KCl. Center: Data and simulations for Hog1-as. Right: Data and simulations for Hog1-as with no salt stimulus. All simulations are n = 1000 and shaded regions are SD = 1. **(E)** Model IIc predictions to previously published data (*English et al., 2015). Left: Data and simulations for wildtype Hog1 in response to 450, 350, 250 mM KCl. Right: Data and simulations for wildtype Hog1-as.

To complete our systematic screen of potential circuitries, we also added positive feedback loops to our Model I to determine whether a positive feedback and direct negative feedback was sufficient to capture our signaling dynamics. Nevertheless, in Models Ia-c, Hog1 did not remain fully phosphorylated, diminishing within the first few minutes (Figure 4 – figure supplement 3). Together, these results support the existence of a delayed negative feedback loop as well as a fast positive feedback loop targeting a component within close proximity of Hog1.

### Experimental validation of computational models reveals positive feedback targeting Hog1

A successful model must not only fit relevant data but also predict new behavior. One particularly informative approach is to use such models to predict the response to dynamic input and determine whether they are able to capture dynamics more complex than those used to train the model. With two pathway circuitries (Models IIb and IIc) that sufficiently captured our data (Figure 5A), we aimed to differentiate them by predicting Hog1 behavior in response to increasing step stimuli. To identify the best model, we sought an input that produced different outputs for each model, and to then test those conditions experimentally (Mélykúti et al., 2010).

**Figure 5:**
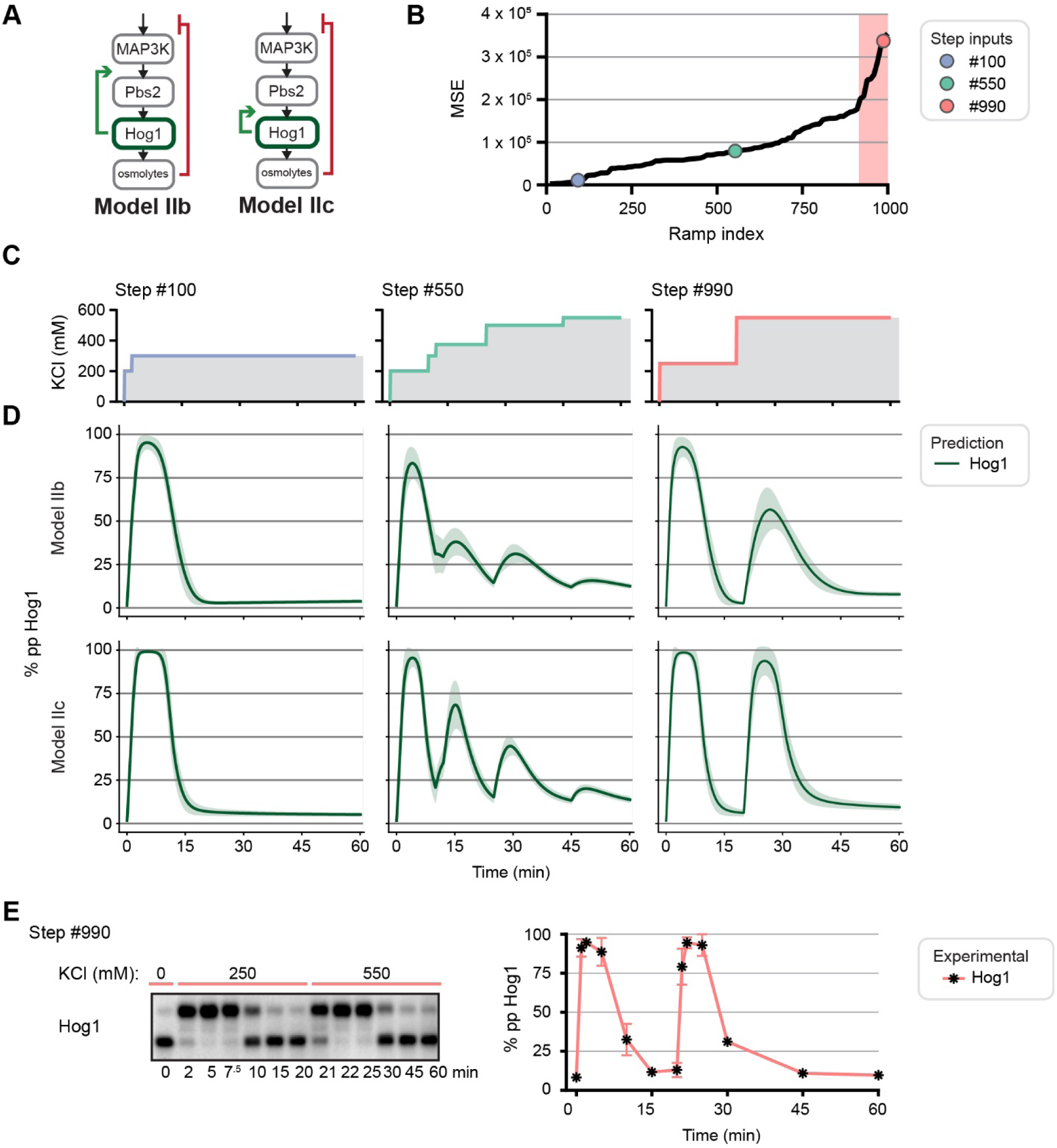
Differentiating models by predicting Hog1 behaviors to dynamic inputs. **(A)** Schematics of the two models that fit our data. **(B)** Mean squared errors (MSEs) for the predicted Hog1 behaviors of Models IIb and IIc for 1000 randomly generated increasing steps. Pink shaded area indicates where step inputs follow a trend similar to that of Steps #990 (pink circle). **(C)** Selected steps depicting a low (left), mid (center), and high (right) scoring step input. **(D)** Predicted Hog1 behaviors to the three step inputs for Models IIb (mid), and IIc (bottom) (C). **(E)** Experimental Hog1 behavior to step stimulus. Left: Hog1 behavior in response to Step #990 resolved using Phos-tag SDS-PAGE (n=3). Right: Quantification of blots. Error bar represent SD of each point.

Following this strategy, we computationally generated 1000 random input profiles of increasing salt concentrations and predicted Hog1 response to each input profile using Models IIb and IIc. These step profiles were designed so that they could be experimentally tested in vivo. We ranked the resulting input profiles based on which generated the largest differences in the Hog1 response (Figure 5B). For example, Figure 5C shows three selected inputs (“Step”) that correspond to the Hog1 dynamics predicted by the two models in Figure 5D. Step #100 generated similar predictions among the models while Step #990 resulted in distinct Hog1 behaviors. Step #550 also predicted model-dependent dynamics, but the differences were too small to be experimentally decipherable. Generally, the input profiles that produced the greatest difference between the Hog1 behaviors were those that allowed Hog1 to adapt to an initial step of KCl before introducing a second step (shaded area in Figure 5B). For Step #990, Model IIb predicted that Hog1 would show a diminished response to the second step of stimulus, but Model IIc predicted full Hog1 phosphorylation in response to this second step (Figure 5D right column). These results indicated that Step #990 would discriminate between the two models.

We then measured the biological Hog1 response to Step #990. We exposed cells to the stimulus profile used in our simulations: beginning with an initial salt stimulus of 250 mM KCl and then raising the salt concentration to 550 mM KCl after 20 minutes. Hog1 activity was again measured by Phos-tag immunoblotting (Figure 5E, left). Quantitation of the blots shows that Hog1 responded normally to the first step of stimulus – becoming completely phosphorylated by two minutes and then fully adapting within 15 minutes (Figure 5E, right). Upon the second stimulus step, Hog1 was again fully activated and then fully adapted. This result was similar to previously published measures of Hog1 translocation and phosphorylation (by phospho-p38 immunoblotting) in response to steps of equal magnitude (Behar et al., 2007; Hao et al., 2007; Zi et al., 2010). In further support of Model IIc, we then predicted Hog1 dual phosphorylation if its kinase activity was inhibited directly before the second stimulus step of Step #990. To conduct this experiment, we utilized the Hog1-as strain. Again, results most closely aligned with Model IIc (Figure 5 – figure supplement 1). Thus, our experimental results to Step #990, both with and without kinase activity, most closely aligned to the predicted Hog1 dynamics of Model IIc, indicating that positive feedback likely acts at the level of Hog1 rather than elsewhere in the MAPK cascade.

### Positive feedback is independent of feedback phosphorylation

Our modeling results suggested that positive feedback amplifies the signal at the level of Hog1. There are two ways in which feedback phosphorylation could activate the MAPK: increase its phosphorylation rate (Figure 6) or decrease its dephosphorylation rate (Figure 7). Since positive feedback must happen quickly, it seemed likely that the target of feedback regulation is a direct substrate of Hog1. Hog1, like all MAPKs, phosphorylates proteins at a serine or threonine followed by a proline. Phosphorylation at these sites typically invokes conformational changes or alters binding affinities, resulting in rapid changes in substrate function (Humphrey et al., 2015; Ubersax and Ferrell, 2007). If positive feedback is due to phosphorylation by Hog1, then mutating the MAPK consensus sites in potential feedback targets should dampen Hog1 activity.

**Figure 6:**
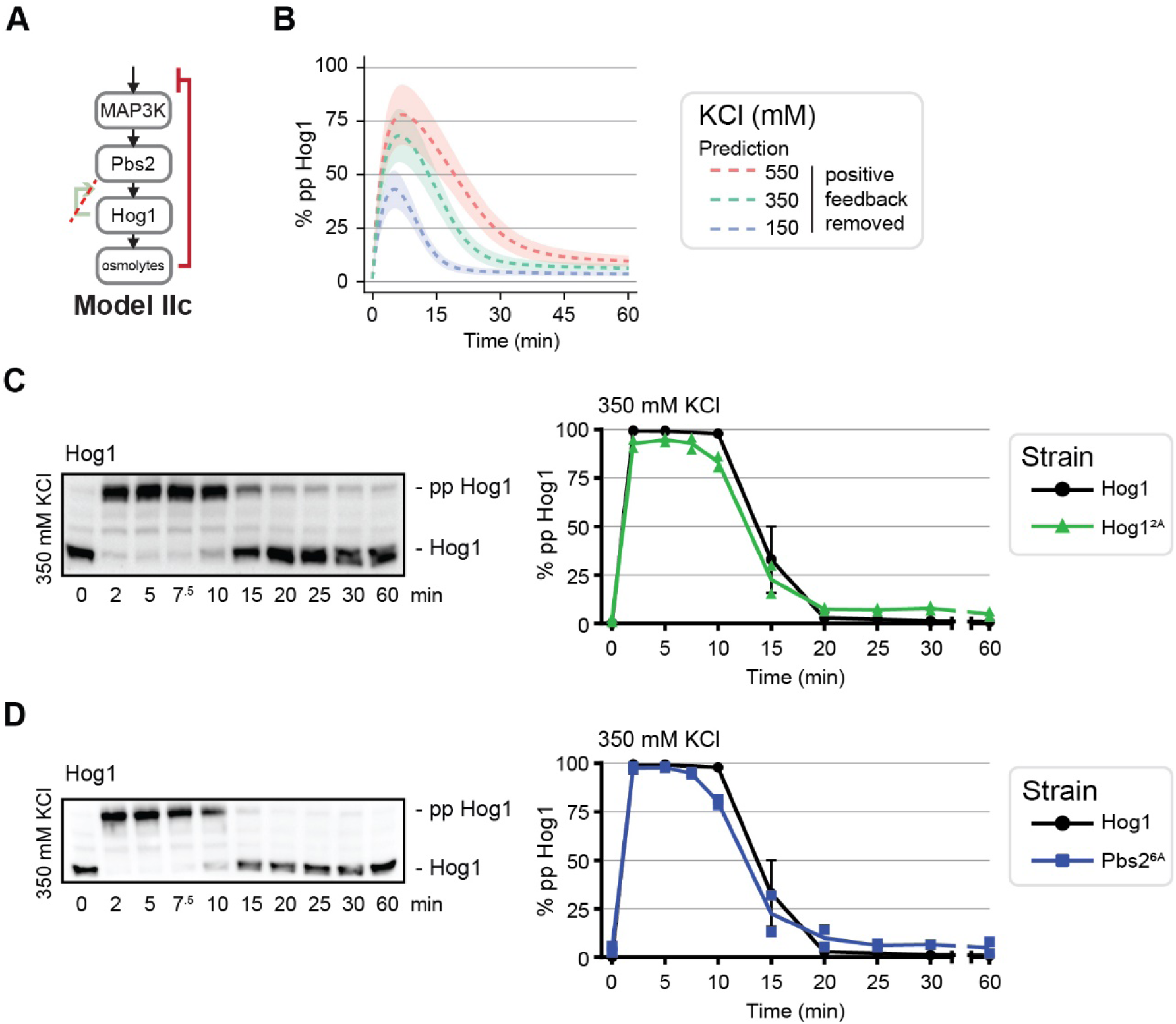
Evaluating increasing Hog1 phosphorylation as the positive feedback mechanism. **(A)** Schematic of Model IIc with positive feedback removed. **(B)** Model IIc prediction of Hog1 in response to 550, 350, and 150 mM KCl without positive feedback. **(C)** Left: Hog1 behavior in response to 350 mM KCl with putative MAPK consensus sites mutated in Pbs2. Right: Quantification of blots. **(D)** Left: Hog1 behavior in response to 350 mM KCl with putative MAPK consensus sites mutated in Hog1. Right: Quantification of blots. n = 2 for mutants, points are replicates and line is mean; n = 3 for wildtype Hog1, points and line are mean with SD.

**Figure 7:**
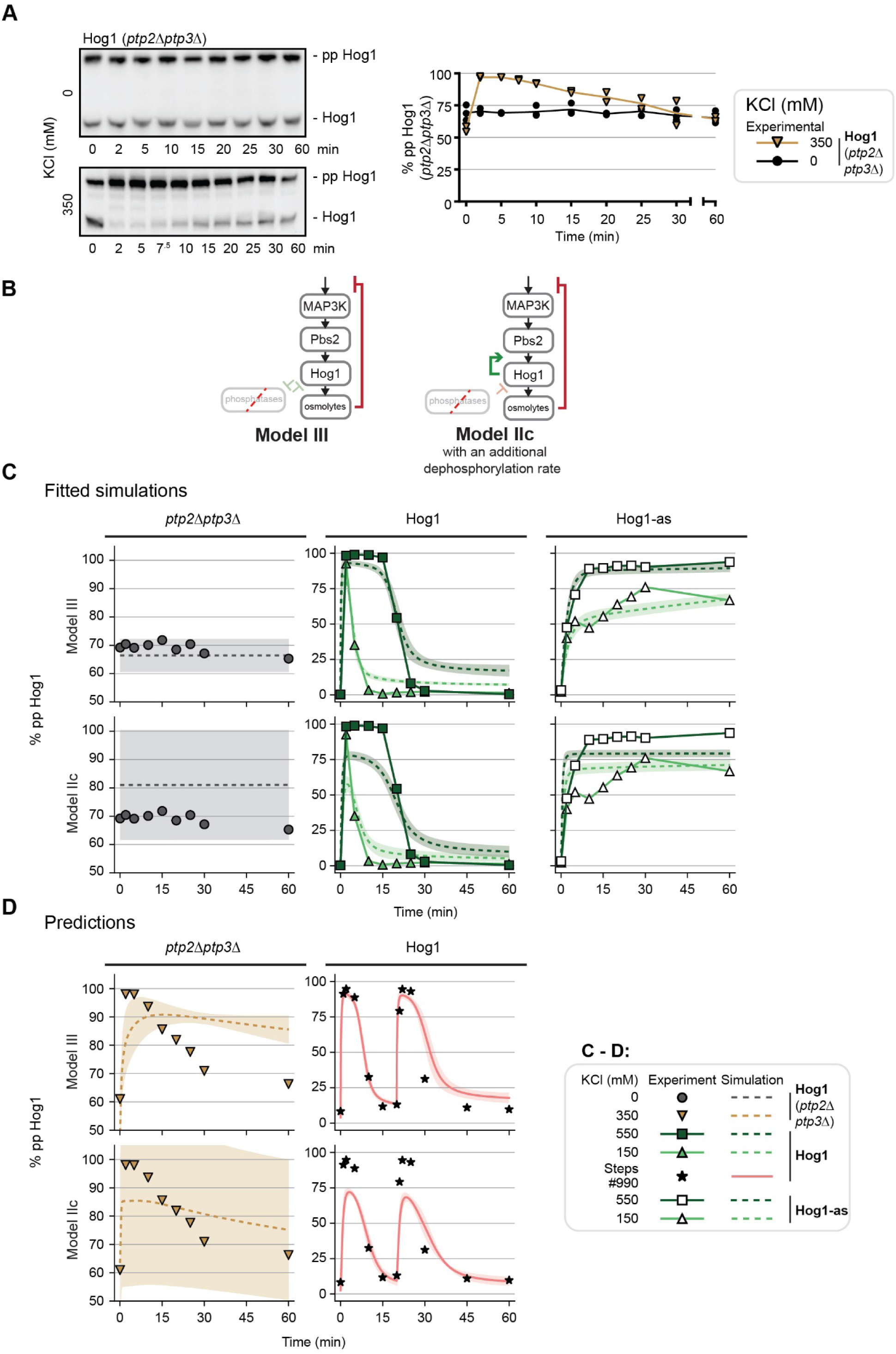
Evaluating decreasing Hog1 dephosphorylation as the positive feedback mechanism. **(A)** Left: Hog1 behavior in response to 350 mM KCl and no KCl in a *ptp2*Δ*ptp3*Δ background. Right: Quantification of blots. n=2, points are replicates and line is mean. **(B)** Schematic of models incorporating phosphatases. Left: Model III, with positive feedback acts through mutual inhibition between Hog1 and its phosphatases. Right: Model IIc, with an additional dephosphorylation rate to simulate the removal of the phosphatases. **(C)** Model fits to experimental data (selective representatives shown). Top: Model III. Left: Simulated Hog1 fits to *ptp2*Δ*ptp3*Δ without KCl stimulus. Center: Fits to wildtype Hog1 dynamics for 550 and 150 mM KCl. Right: Fits to Hog1-as data. Bottom: Same as Top but for Model IIc. **(D)** Model predictions compared to experimental data. Top: Model III. Left: Prediction of Hog1 dynamics in response to 350 mM KCl in a *ptp2*Δ*ptp3*Δ background. Right: Prediction in response to Steps #990. Bottom: Same as Top but for Model IIc.

We then used Model IIc to investigate how feedback phosphorylation could amplify Hog1 phosphorylation. By assigning the activation rate α to 0, thereby eliminating positive feedback, the model predicted a reduction in Hog1 phosphorylation, particularly at low salt concentrations (Figure 6B). Based on these predictions, we anticipated that 350 mM KCl would be particularly informative since it was low enough to cause at least a 25% decrease in Hog1 phosphorylation over several timepoints (Figure 6B). To disrupt the putative positive feedback loop, we mutated the two MAPK consensus sites on Hog1 (Hog1^2A^ mutant) and monitored its phosphorylation in response to 350 mM KCl. Immunoblotting after Phos-tag SDS-PAGE showed that these mutations did not alter Hog1 dynamics (Figure 6C), in contrast to predictions of Model IIc. We then considered Pbs2 as a potential substrate since it is responsible for Hog1 activation. We mutated its 6 MAPK consensus sites (Pbs2^6A^ mutant), and found that these alterations also produced minimal changes in Hog1 activation (Figure 6D). Taken together, these results suggest that phosphorylation of Pbs2 or Hog1 is not the source of positive feedback in the system.

### Positive feedback results from mutual inhibition of Hog1 and its phosphatases

We then considered an alternative scenario where Hog1 acts by decreasing its own rate of deactivation. In practice, this could be achieved by Hog1 inhibition of its phosphatases. We constructed a new model, Model III, that incorporated another model species representing Hog1 phosphatases and included mutual inhibition between the phosphatases and Hog1.

*Model III*:

consisted of Model II’s equations 1, 2, 3, and 4 with the following modifications:

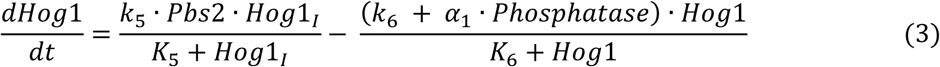

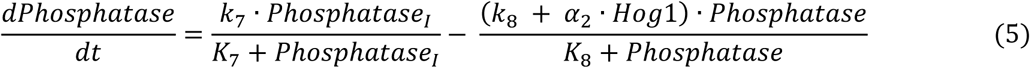

where *α*_1_ is phosphatase-driven Hog1 suppression and *α*_2_ is Hog1-driven phosphatase suppression. Here, the total phosphatase concentration is conserved.

We determined whether Model III could perform equal to or outperform Model IIc. We trained Model III on the same phosphorylation data for both Pbs2 and Hog1, as previously done for Model IIc (Figure 3). Resulting fits to Model III captured our 10 pathway characteristics as well as Model IIc (Figure 7 – figure supplement 1). Based on these results, we conclude that mutual inhibition is a candidate for positive feedback in HOG signaling pathway.

Next, we sought to gain experimental evidence in support of our mutual inhibition hypothesis. We examined three potential targets of mutual inhibition: the Hog1-directed phosphatases Ptc1, Ptp2, and Ptp3 (Jacoby et al., 1997; Mattison and Ota, 2000; Warmka et al., 2001; Wurgler-Murphy et al., 1997). Ptc1 dephosphorylates Hog1 at its activation loop threonine site while Ptp2 and Ptp3 dephosphorylate the remaining tyrosine site. Among these three phosphatases there are 22 putative MAPK consensus sites. Hog1 could phosphorylate a combination of these sites to suppress phosphatase activity. Since mutating every site was infeasible, we instead deleted the *PTC1, PTP2, PTP3* genes and monitored Hog1 phosphorylation. Each deletion caused mild changes to the timing of Hog1 dephosphorylation, but did not result in the partial Hog1 activation that the model predicted (Figure 7 – figure supplement 2). This result suggests that a single phosphatase is unlikely to be responsible for feedback regulation.

Existing evidence indicates that the three phosphatases work together to regulate Hog1, making it likely that Hog1, in turn, inhibits multiple phosphatases. In particular, dual deletions of *PTC1* and *PTP2* are lethal, most likely due to Hog1 hyperactivation (Maeda et al., 1993). Our previously published data showed that Hog1 was basally phosphorylated in a *ptp2*Δ*ptp3*Δ background (English et al., 2015). Additional investigation revealed that deletion of both *PTP2* and *PTP3* results in high (70%) basal phosphorylation of Hog1 (Figure 7A); in response to 350 mM KCl, Hog1 is fully phosphorylated and then returns back to 70% basal activation. Though this experimental result alone is insufficient to suggest that positive feedback acts through mutual inhibition, we could nevertheless use this data to retrain our models to determine if positive feedback was needed in the system.

To distinguish between the mutual inhibition and positive feedback loop mechanisms, we compared how well Model III and IIc fit our *ptp2*Δ*ptp3*Δ data. If this mutual inhibition applied, Model III should be able to capture all of the phosphorylation data, indicating that positive feedback is not present within a *ptp2*Δ*ptp3*Δ background. In contrast, should Model IIc capture this data, this would imply positive feedback is still present within a *ptp2*Δ*ptp3*Δ background, since we are only eliminating phosphatase suppression of Hog1 activity but not the positive feedback loop. Thus, we retrained Model IIc and III to the basal phosphorylation of the *ptp2*Δ*ptp3*Δ data and compared their performance. For Model IIc, we simulated *ptp2*Δ*ptp3*Δ by fitting a separate Hog1 deactivation rate. For Model III, we simulated phosphatase deletion by setting their concentration to 0 (Figure 7B). Fitting to these additional data, we found that Model III was able to capture the experimental data (Figure 7C, top) whereas Model IIc could not, particularly in the wildtype strain (Figure 7C, bottom). Model III could also correctly predict the Hog1 response to Step #990 (Figure 7D, top right) and nearly predicted the behavior of the *ptp2*Δ*ptp3*Δ strain to a single step of 350 mM KCl (Figure 7D, top left). The only discrepancy between Model III and the experimental result was that the experimental measurements for the *ptp2*Δ*ptp3*Δ strain showed faster adaptation than predicted in our simulations. However, this faster dephosphorylation is likely driven by other yeast phosphatases not present in the model. Meanwhile, the retrained Model IIc poorly predicted the Hog1 dynamics in response to 350 mM KCl (Figure 7D, bottom left) and the full Hog1 phosphorylation in response to Step #990 (Figure 7D, bottom right). The performance of Model III, in both its fits to the data and its prediction of the increasing step stimulus behavior, provides strong evidence for mutual inhibition between Hog1 and its phosphatases. We conclude that mutual inhibition is responsible for positive feedback in the HOG MAPK cascade.

## Discussion

Feedback regulation often controls the timing of signaling events, allowing for an appropriate cellular response. For the HOG pathway, we and others have previously shown that a progressively stronger input leads to a progressively longer output (Aymoz et al., 2016; Behar et al., 2008; English et al., 2015). What has been lacking is a comprehensive understanding of the feedback mechanisms responsible for the encoding of this distinctive “dose-to-duration” signaling profile. To elucidate these mechanisms, we systematically tested 8 network architectures and found two that could fit our experimental data. By changing the input profile and predicting Hog1 response, we found conditions that could differentiate between the two models. Experimental validation identified slow negative feedback and fast positive feedback as the most likely circuitry. We then tested potential mechanisms of positive feedback, with our data suggesting positive feedback acts through mutual inhibition between Hog1 and the tyrosine phosphatases, Ptp2 and Ptp3. Thus, our iterative approach allowed us to identify new mechanisms of regulation in the canonical HOG pathway.

Our findings build on other investigations of feedback within the HOG pathway. Our own previous models incorporated positive feedback, but did not explore how positive feedback acts in conjunction with negative feedback to control Hog1 activation dynamics (English et al., 2015). The present work highlights the importance of tyrosine phosphatases together with osmolyte accumulation. However, other feedback mechanisms are likely to be important for controlling Hog1 dynamics. These could include known mechanisms, such as Hog1 phosphorylation of upstream components and other Hog1-directed phosphatases. Other mechanisms of feedback have been suggested, particularly between the two input branches which seem to suppress one another’s activity (Granados et al., 2017). Thus, feedback likely acts on a variety of components to continuously fine-tune the cell’s response to a given stimulus. Looking forward, investigating the response of Hog1 to even more complex inputs, including different ramps (Thiemicke et al., 2019) or pulses, will further clarify the roles of individual feedbacks within the system.

More generally, the results provided here suggest that the counter-acting mechanisms of positive and negative feedback determine the prioritization of intracellular events following hyperosmotic stress. These events are likely to occur on various timescales. For example, shortly after the stimulus, Hog1 phosphorylates a regulator of Fps1, a glycerol export channel, resulting in rapid channel closure and the accumulation of glycerol in the cell (Lee et al., 2013). On a longer timescale, Hog1 phosphorylates transcription factors resulting in new gene expression (Alepuz et al., 2001; Capaldi et al., 2008). With prolonged stimulation, Hog1 activates multiple transcription factors and in so doing employs additional regulatory mechanisms such as feedforward loops (AkhavanAghdam et al., 2016). The timing of these actions suggests a prioritized order of intracellular events, presumably to enhance a cell’s chance of surviving hyperosmotic stresses.

Collectively, these efforts illustrate how computational modeling allows us to probe behaviors that are difficult to predict or explain through experimentation alone. When models are based on quantitative data and describe well-defined molecular networks, it is possible to extract information about the system and make predictions of how that system behaves under complex situations. Here we found step stimuli that could differentiate the predicted behaviors of models that captured our experimental data. This model-driven experimental design not only provided insights into circuit-specific behaviors, but it also revealed putative mechanisms of positive feedback.

Likewise, insights developed from the yeast system could reveal regulatory roles of other MAPKs in more complex systems. In a broader context, understanding how pathways control MAPK regulation is critical for pharmaceutical development. Protein kinases are the second largest group of drug targets, and are particularly important in the treatment of cancers. Moreover, one of the main challenges of drug development is overcoming kinase inhibitor resistance within complex pathway systems (Bhullar et al., 2018). Understanding the mechanisms of spatiotemporal pathway regulation will ultimately lead to the development of novel techniques to control kinase activity.

## Materials and Methods

### Strain construction and plasmids

Strains (Table 1) were derived from BY4741 (“wildtype”) and transformed by the lithium acetate method (Gietz and Woods, 2002). Pbs2-9xMyc-tagged strains were generated by homologous recombination of a PCR-amplified 9xMyc cassette at the C-terminus of the *PBS2* open reading frame. This cassette contained a resistance gene to hygromycin from plasmid pYM20 (pYM20-9xMyc-hphNT1) (Janke et al., 2004).

**Table 1.**
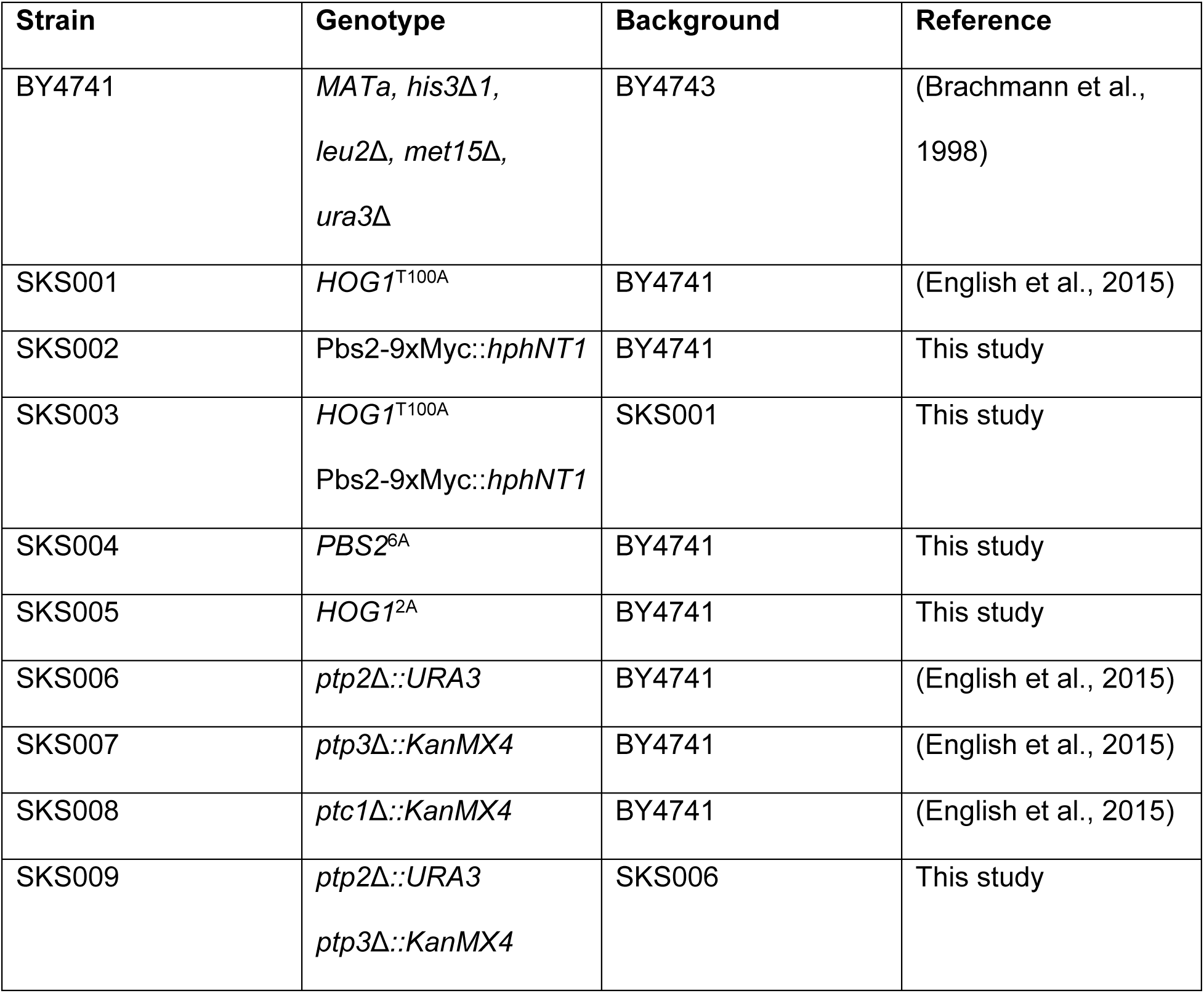

| Strain | Genotype | Background | Reference |
| --- | --- | --- | --- |
| BY4741 | *MATa, his3Δ1, leu2Δ, met15Δ, ura3Δ* | BY4743 | (Brachmann et al., 1998) |
| SKS001 | *HOG1<sup>T100A</sup>* | BY4741 | (English et al., 2015) |
| SKS002 | *Pbs2-9xMyc::hphNT1* | BY4741 | This study |
| SKS003 | *HOG1<sup>T100A</sup>*<br>*Pbs2-9xMyc::hphNT1* | SKS001 | This study |
| SKS004 | *PBS2<sup>6A</sup>* | BY4741 | This study |
| SKS005 | *HOG1<sup>2A</sup>* | BY4741 | This study |
| SKS006 | *ptp2Δ::URA3* | BY4741 | (English et al., 2015) |
| SKS007 | *ptp3Δ::KanMX4* | BY4741 | (English et al., 2015) |
| SKS008 | *ptc1Δ::KanMX4* | BY4741 | (English et al., 2015) |
| SKS009 | *ptp2Δ::URA3*<br>*ptp3Δ::KanMX4* | SKS006 | This study |

Mutagenesis for Hog1 (S91A and S235A) and Pbs2 (S83A, T164A, T212, S248A, T297A, and S415A) were introduced using the *delitto perfetto* method (Stuckey et al., 2011) using the PCR-amplified pCORE cassette (RRID:Addgene_72231) to integrate selective markers at the endogenous gene loci. These markers were selected against after the integration of synthesized gBlocks (Integrated DNA Technologies). All strains were validated with PCR, and mutated genes were PCR-amplified and sequenced.

### Cell culture

Strains were cultured using standard methods and media. Strains were struck out on YPD (yeast extract, peptone, and 2% dextrose) plates and cultured at 30°C. Individual colonies were picked and grown overnight in 3 mLs SCD (synthetic complete and 2% dextrose) medium to saturation. Cells were diluted 1:100, grown for 8 hours, and diluted to OD_600_ = 0.001 for overnight growth. The following day, experiments were conducted once the cell culture reached an OD_600_ ∼1.

### Phos-tag sample collection, gel electrophoresis, and immunoblotting

Kinase activation was quantified using Phos-tag immunoblotting technique as previously described (English et al., 2015). Briefly, cells were cultured with a final volume of 80 mLs in SCD. For Hog1-as (Hog1^T100A^ + 1-NA-PP1) kinase inhibition, 1-NA-PP1 ATP analog (Cayman Chemical, #10954) was added to cultures to a final concentration of 12 µM and incubated for 2 min before sampling. At the selected timepoints after the addition of KCl in SCD, samples were quenched in 5% (v/v) trichloroacetic acid (TCA) on ice, washed with 5% sodium azide, and stored at −80°C. Sample concentrations were normalized to 1.5µg/µL using the DC Protein Assay (Bio-Rad) and stored at −80°C.

Samples were resolved using 8% acrylamide 20uM Phos-tag Bis-Tris SDS-PAGE gels and transferred on to PVDF membrane. Hog1 was detected using an anti-Hog1 primary-antibody (Santa Cruz, Hog1 antibody (D-3) sc-165978; 1:5,000) and a donkey-anti mouse HRP-conjugated secondary antibody (Jackson ImmunoResearch, 715-035-150; 1:10,000). Pbs2-9xMyc was detected using an anti-Myc primary antibody (Cell Signaling, 9B11 #2276, 1:5,000) and a donkey anti-rabbit HRP-conjugated secondary antibody (Jackson ImmunoResearch, 711-035-152; 1:10,000). Secondary antibodies were visualized using Clarity Western ECL Substrate (Bio-Rad, #1705061) and a BioRad Chemidoc Touch Imaging System. Band intensities were normalized and quantified using the ImageLab (Bio-Rad) software. We found that additional bands were occasionally observed, that would vary between technical replicates, indicating that their existence was due to gel and immunoblotting inconsistencies rather than being other phospho-states of Hog1. Also, band migration depended on the number of gels run simultaneously. Standard error of the mean was plotted since models were fit to mean values.

### Glycerol measurements

Samples of 1 mL were collected at the selected timepoints after the addition of KCl in SCD and kinase inhibition, when applicable, as above. 500 µL was used to measure OD_600_ and the remaining 500 µL was pelleted and frozen in liquid nitrogen. After collection, samples were boiled for 10 min in sterile water and cleared by centrifugation. The concentration of glycerol was measured using a Free Glycerol Assay Kit (abcam, ab65337) following the manufacturing instructions. Conversion between OD_600_ and cell number was calculated by counting the cells growing in liquid culture with a hemocytometer and measuring the OD_600_ simultaneously (n = 3). These measurements were fit using logarithmic function, which served as a standard curve for our sample measurements to calculate cell number.

### ODE modeling and parameter optimization

Modeling was performed in Python 3.7 using the scipy package to solve ODE systems and their steady states. All kinases and phosphatases observed mass conservation with the total protein amounts reflecting biologically observed concentrations (Ho et al., 2018). These models rely on different assumptions. First, we do not include synthesis or degradation of the kinases because hyperosmotic stress does not induce their transcription (O’Rourke and Herskowitz, 2004) and quantification of Hog1 and Pbs2 time course immunoblots indicates that protein concentration does not change appreciably throughout our experiments (data not shown). Furthermore, we group the three HOG pathway MAP3Ks into one species, assuming that they share the same kinetic behavior. We reason that we are studying the overall behavior of Pbs2 and Hog1, which are downstream of the two input branches.

For parameter optimization, we combined two approaches that have been used to parameterize ODE models to experimental data: an evolutionary algorithm (EA) (Fortin et al., 2012) and an approximate Bayesian Computation and sequential Monte Carlo (ABC SMC) (Toni et al., 2009). All values for *k*_cat_, K_M_, synthesis, degradation, and feedback terms needed to be estimated to fit each model to our experimental data.

First, the EA seeded each simulation with starting values that were randomly selected from a user specified range. Then, the EA would evaluate the fits of each parameter set to the experimental data using MSE and select the best fitting parameter sets to continue to the next generation. To avoid local optima, each parameter set has a 10% probability to crossover with another set, and each parameter has a 20% probability to mutate to a different value. For each model, we calculated the fit of 500 parameter sets over 1000 generations for 2000 independent runs. For each run, we saved the top fitting parameter set. We noticed that it was difficult to programmatically separate out the top fitting parameter sets: when we ranked the MSEs, there was a sharp increase in MSEs, then a gradual increase, followed by another sharp increase. Where these transitions occurred varied with each model, and their resulting fits to the data also depended on the model.

Thus, we chose to use the best (lowest-scoring) 500 EA parameters vectors from the EA as priors for the ABC SMC to further sample for the optimal parameters of each model. This loose inclusion of the best 25% parameter sets allowed the ABC SMC to further search the parameter space in case the EA missed any optima. We then followed the same algorithm as in Toni 2009 in which sampled parameter vectors must pass a series of tolerance levels (ϵ) determined by their fit to the experimental data, where the first tolerance was the worst MSE of the top 25% EA parameter sets and all subsequent tolerances were the average of the previous tolerance and the best MSE from the top 25% EA parameter sets. For each model, we ran four series, or “schedules,” in which each schedule included 1000 parameter vectors that passed its tolerance. During a schedule, a parameter vector was selected based on its importance weight and perturbed. This weight is calculated by the prior and the perturbation of each parameter. We used a perturbation kernel of U(−1,1) around log_10_ transformed parameter values so that sampling was scaled to the magnitude of the value. Since all priors and perturbation kernels for these simulations were uniform, each parameter set had an equal probability of being selected. After each schedule, we calculated new weights for the selected parameter values. In the end, we had 1000 parameter vectors that passed the highest tolerance threshold.

All simulations and analysis were performed using custom scripts which are available at https://github.com/sksuzuki/HOG_encoding_feedbacks.

### Model differentiation

Once we found two models that could capture our experimental data, we needed to identify the most likely circuitry of the two. We generated increasing step stimuli and simulated Hog1 response with each model. Each stimulus was randomly generated, but we limited them to three rules. First, the stimulus must always increase because decreasing osmolarity would activate a hypoosmotic response. While the Sln1 branch contributes to the hypoosmotic response, there are other mechanisms outside of the HOG pathway that control yeast response to hypoosmotic stress (Brown et al., 1994). Second, we limited the increasing steps to a maximal stimulus of 550 mM KCl due to increased cell death above this concentration. Third, we limited the intervals of each step to at least 2 minutes since faster intervals are not experimentally feasible. Generated inputs were then ranked based on maximizing the distance between model predictions. Thus, the larger distance reflected the greatest difference between the simulated Hog1 dynamics.

## Author Contributions

SKS, BE, HGD, and TCE were responsible for conceptualization. SKS, HGD, and TCE designed experiments. SKS constructed strains and performed experiments, data analysis, and mathematical modeling. SKS created visualizations. SKS, HGD, and TCE were responsible for writing with review assistance from BE.

## Acknowledgments

We thank Jeremy Purvis, Joel Parker, Michael Pablo, and Amy Pomeroy for helpful discussion, James Shellhammer for guidance on the experimental procedures, Jeff Snell for assistance in developing the evolutionary algorithm, Rozemarijn van der Veen for editing the manuscript, and Joseph McGirr for feedback. This work was supported by the National Institutes of Health (NIH) training grant T32 GM067553-12 (SKS), the NIH Maximizing Investigators’ Research Award (MIRA) R35 GM127145 (TCE), and the NIH MIRA R35 GM118105 (HGD).

---

**Figure 3 – figure supplement 1:**
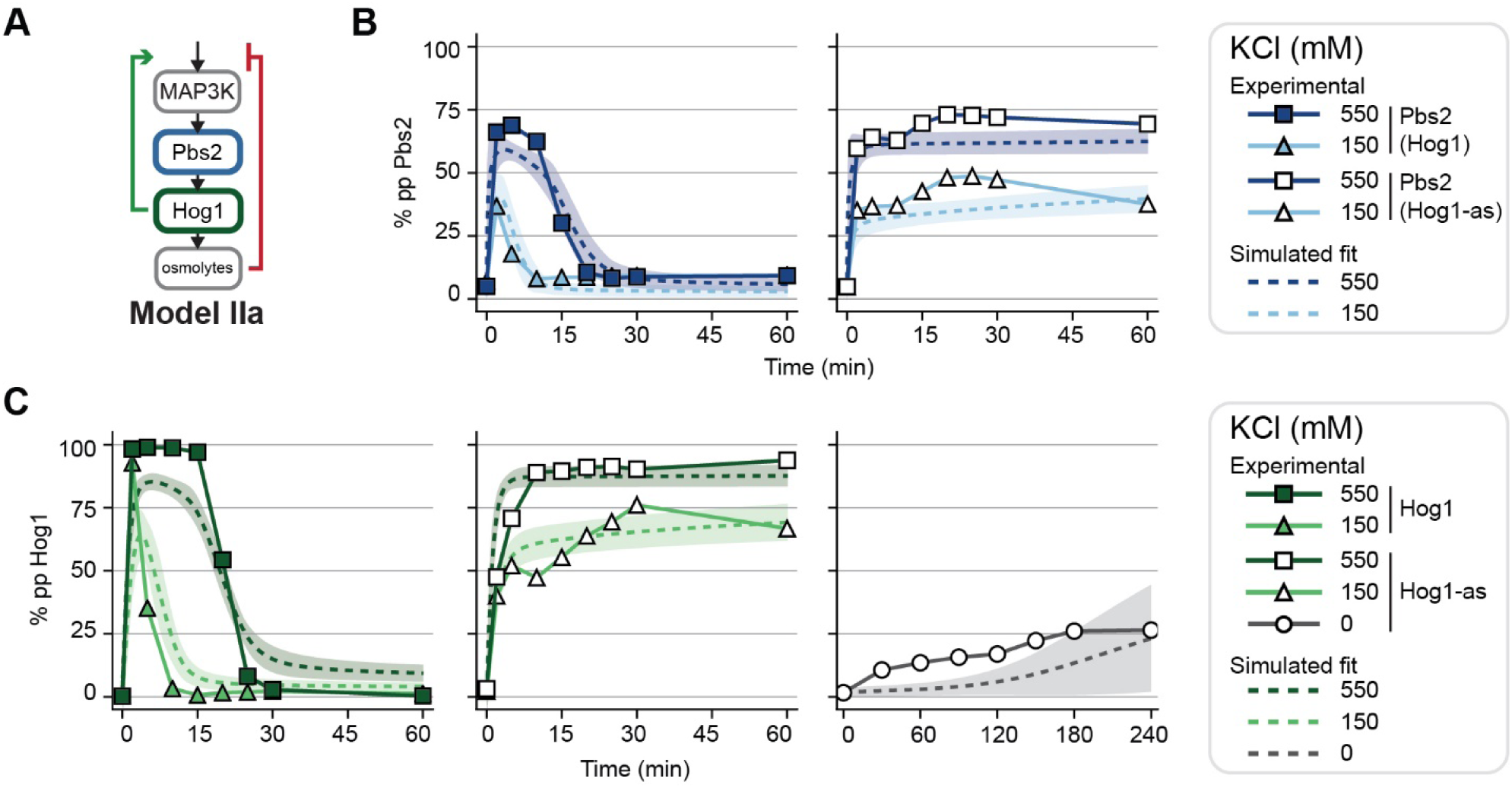
Model II (delayed negative feedback) fits to experimental data. **(A)** Schematic of Model II with negative feedback driven by a species downstream of Hog1, such as Hog1-dependent accumulation of osmolytes. **(B)** Model II simulated Pbs2 fits (dashed lines) overlaid with data (symbols). Left: Data and simulations for wildtype Hog1 in response to 550 mM and 150 mM KCl. Right: Data and simulations for Hog1-as. **(C)** Model II simulated Hog1 fits (dashed lines) overlaid with data (symbols). Left: Data and simulations for wildtype Hog1 in response to 550 mM and 150 mM KCl. Center: Data and simulations for Hog1-as. Right: Data and simulations for Hog1-as with no salt stimulus. All simulations in (B) and (C) are n = 1000 and shaded regions are SD of 1.

**Figure 3 – figure supplement 2:**
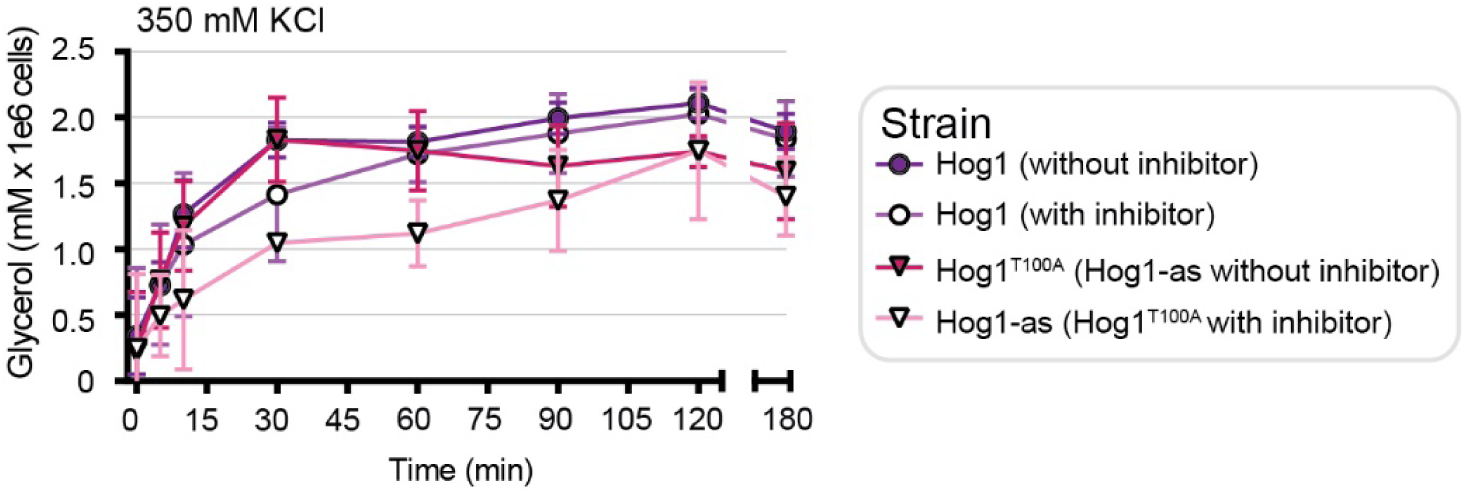
Inhibitor- and Hog1 analog sensitive variant-dependent glycerol accumulation in response to hyperosmotic stress. Glycerol accumulation over time in response to 350 mM KCl with and without 1-NA-PP1 drug in both wildtype Hog1 and Hog1^T100A^ backgrounds. All experiments are n = 3 and error bars represent SD of each point.

**Figure 3 – figure supplement 3:**
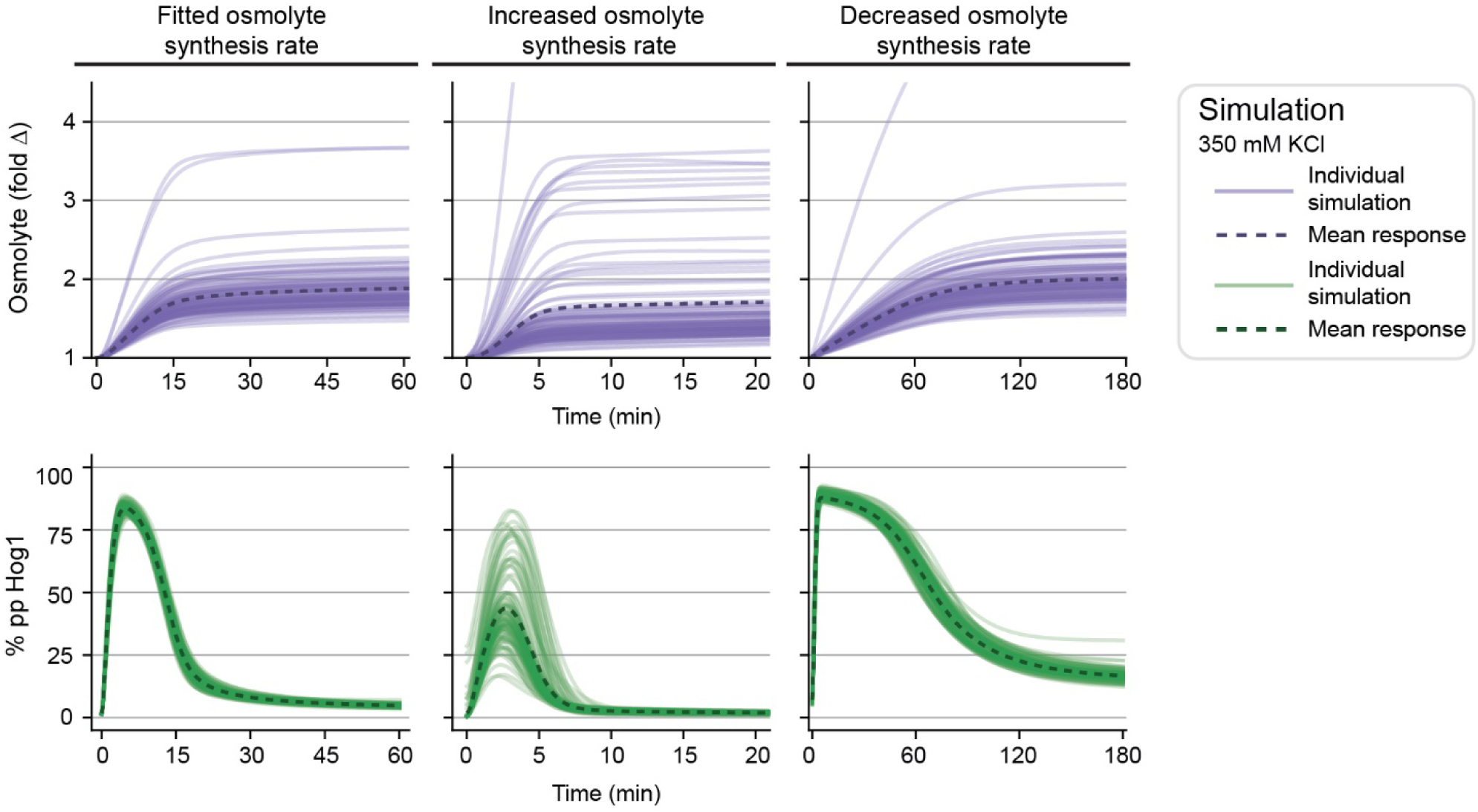
Delayed feedback investigation. Comparing Model II simulations with the fitted (left) osmolyte synthesis rate to 5x increased (center) or 5x decreased (right) osmolyte synthesis rate in response to 350 mM KCl. Each solid line is one simulation corresponding to one fitted parameter set and each dashed line is the mean response of the plotted simulations. Top row is osmolyte simulations (purple) and bottom row is the corresponding Hog1 simulations (green). The best 100 simulations are plotted for clear visualization.

**Figure 4 – figure supplement 1:**
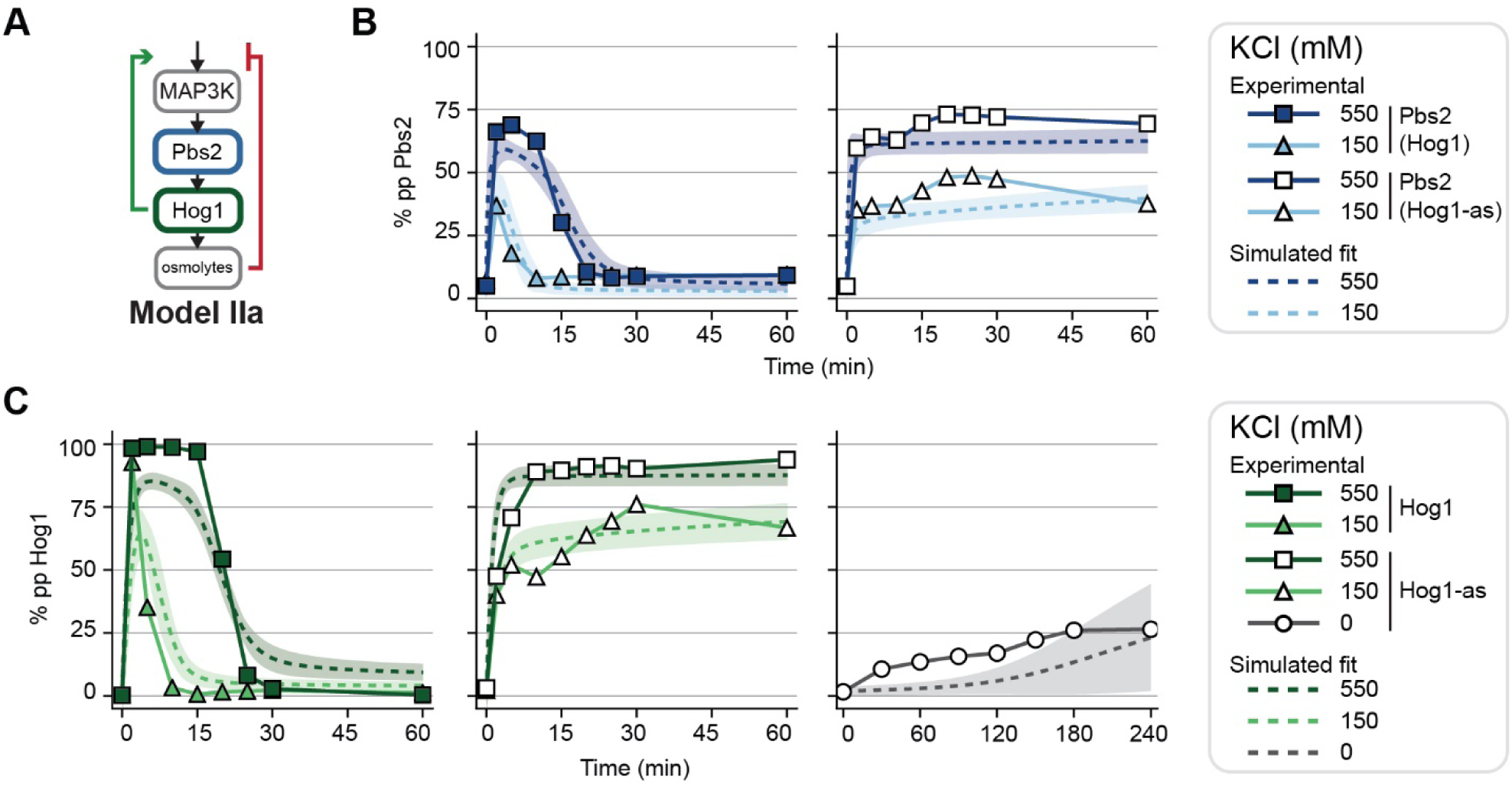
Model IIa with positive and negative feedback poorly fits experimental data. **(A)** Schematic of Model IIa with a delayed negative feedback and a positive feedback increasing MAP3K activation. **(B)** Model IIa simulated Pbs2 fits (dashed lines) overlaid with data (symbols). Left: Data and simulations for wildtype Hog1 in response to 550 mM and 150 mM KCl. Right: Data and simulations for Hog1-as. **(C)** Model IIa simulated Hog1 fits (dashed lines) overlaid with data (symbols). Left: Data and simulations for wildtype Hog1 in response to 550 mM and 150 mM KCl. Center: Data and simulations for Hog1-as. Right: Data and simulations for Hog1-as with no salt stimulus.

**Figure 4 – figure supplement 2:**
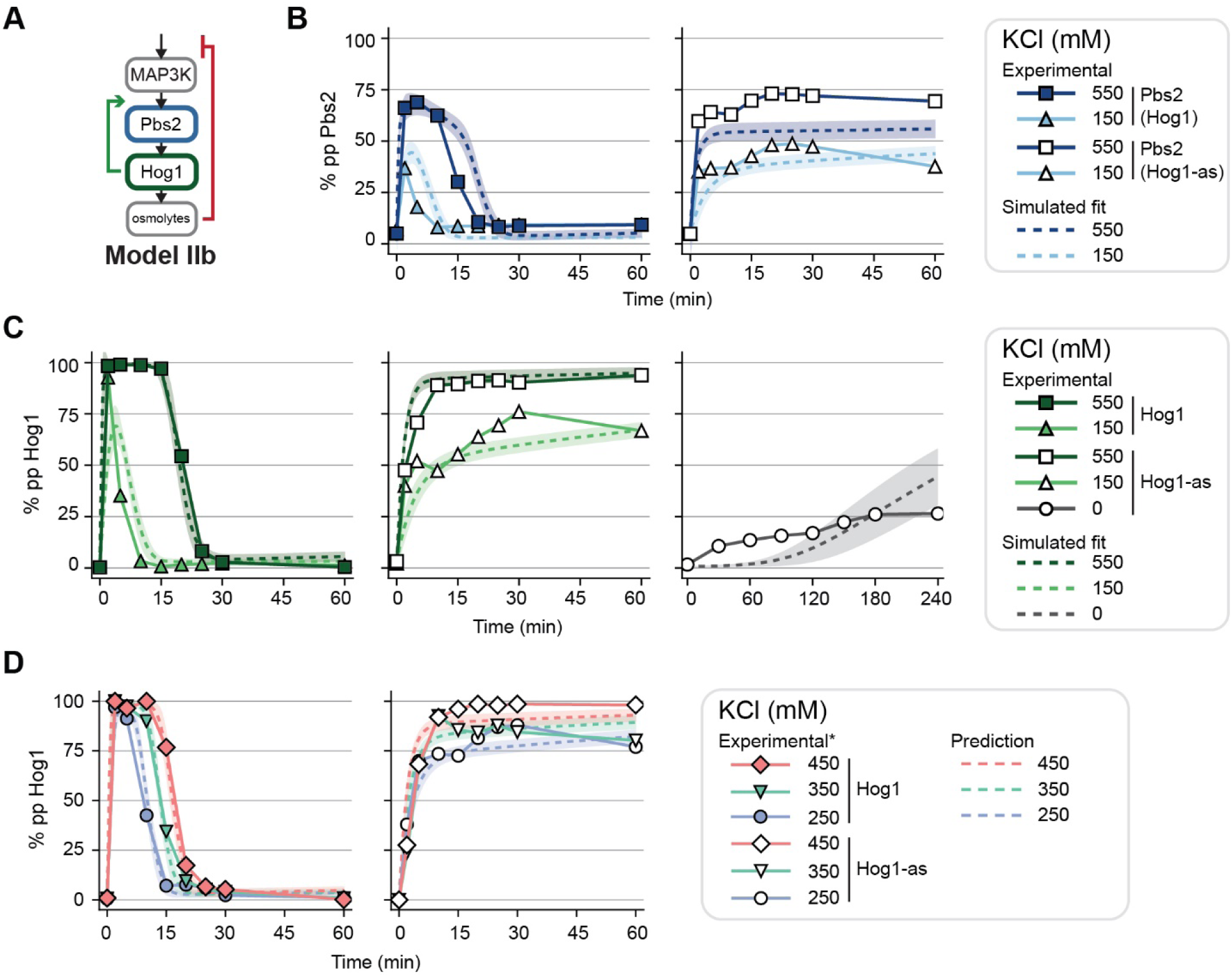
Model IIb with positive and negative feedback that captures experimental data. **(A)** Schematic of Model IIb with a delayed negative feedback and a positive feedback increasing MAP2K activation. **(B)** Model IIa simulated Pbs2 fits (dashed lines) overlaid with data (symbols). Left: Data and simulations for wildtype Hog1 in response to 550 mM and 150 mM KCl. Right: Data and simulations for Hog1-as. **(C)** Model IIa simulated Hog1 fits (dashed lines) overlaid with data (symbols). Left: Data and simulations for wildtype Hog1 in response to 550 mM and 150 mM KCl. Center: Data and simulations for Hog1-as. Right: Data and simulations for Hog1-as with no salt stimulus. **(D)** Model IIb predictions to *previously published data (English et al., 2015). Left: Data and simulations for wildtype Hog1 in response to 450, 350, 250 mM KCl. Right: Data and simulations for wildtype Hog1-as. All simulations are n = 1000 and shaded regions represent a SD of 1.

**Figure 4 – figure supplement 3:**
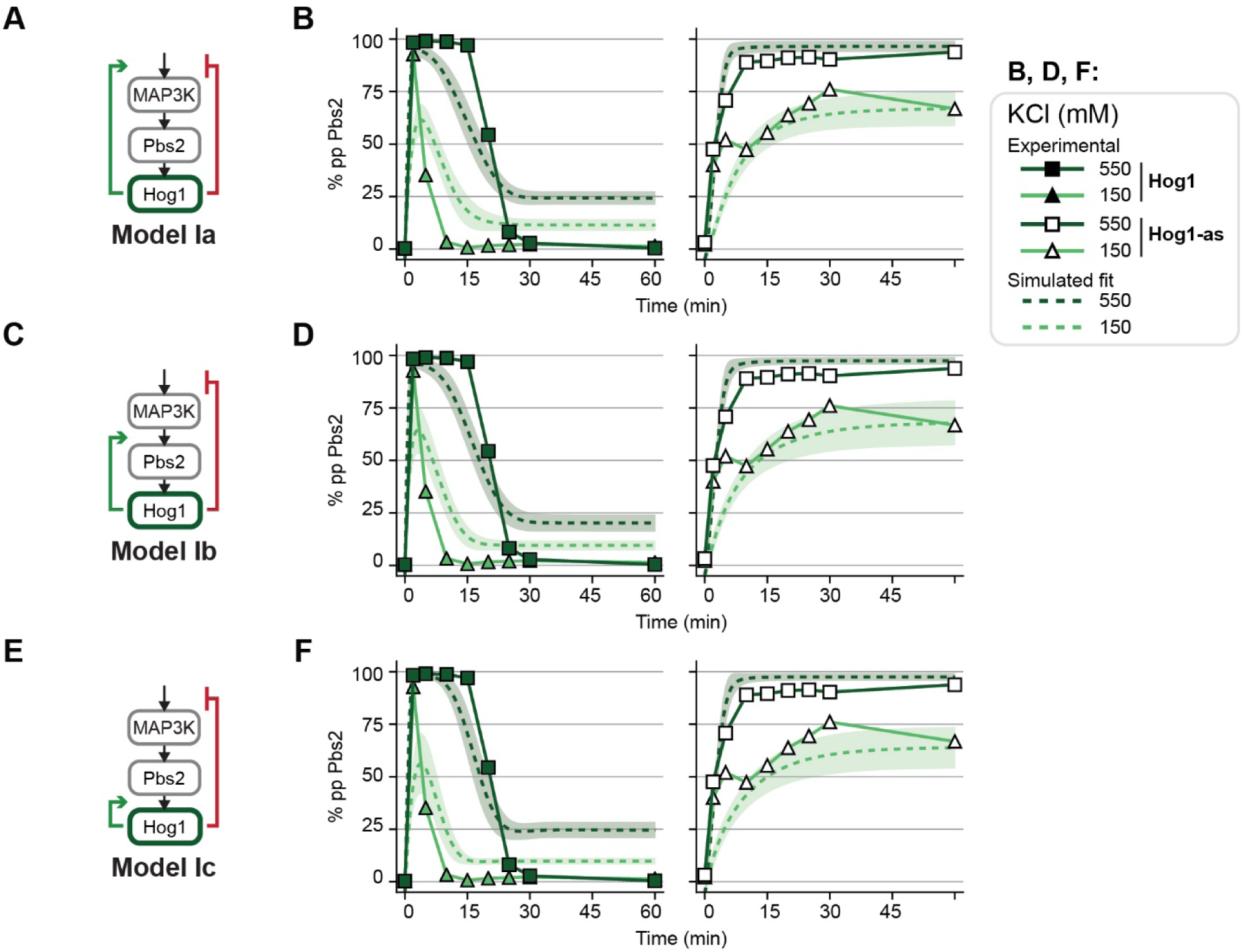
Models with direct negative feedback and positive feedback cannot capture experimental data. **(A)** Schematic of Model Ia with a negative feedback directly from Hog1 and a positive feedback increasing MAP3K activation. **(B)** Model Ia simulated Hog1 fits overlaid with data. Data and simulations for wildtype Hog1 in response to 550 mM and 150 mM KCl. Right: Data and simulations for Hog1-as. **(C)** Schematic of Model Ib with a negative feedback directly from Hog1 and a positive feedback increasing MAP2K activation. **(D)** Same as (B) but for Model Ib. **(E)** Schematic of Model Ic with a negative feedback directly from Hog1 and a positive feedback increasing MAPK activation. **(F)** Same as (B) but for Model Ic. All simulations are n = 1000 and shaded regions represent a SD of 1.

**Figure 5 – figure supplement 1:**
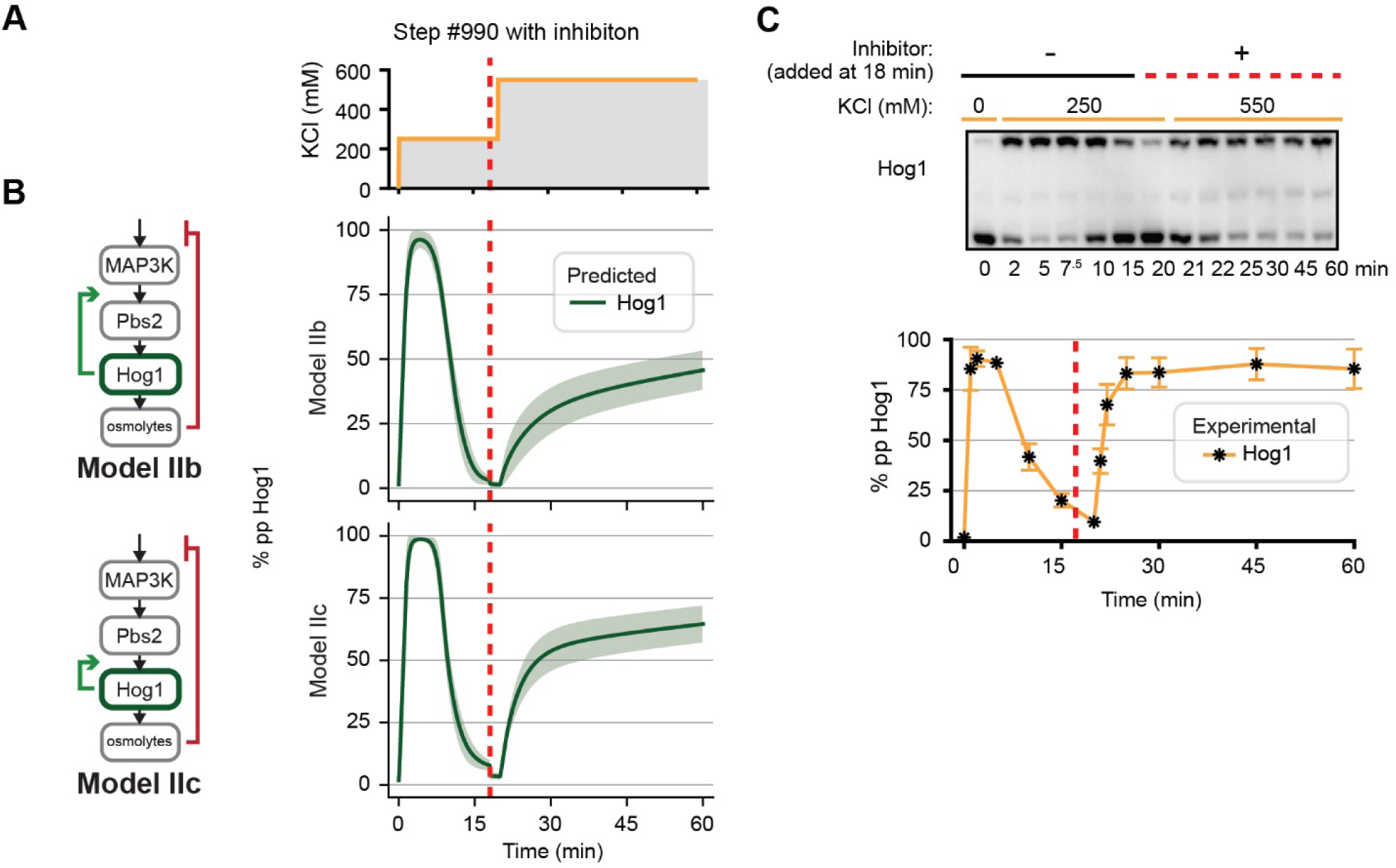
Model and experimental Hog1 behavior in response to step stimulus and inhibition. **(A)** Step stimulus #990 with inhibition before the second step of inhibition (t=18 min). **(B)** Model predictions to step stimulus with the inhibition. Top: Model IIb prediction. Bottom: Model IIc prediction. **(C)** Experimental Hog1 behavior to the stimulus. Top: Hog1 behavior in response to Step #990 resolved using Phos-tag SDS-PAGE (n=3). Bottom: Quantification of blots. Error bar represent SD of each point.

**Figure 7 – figure supplement 1:**
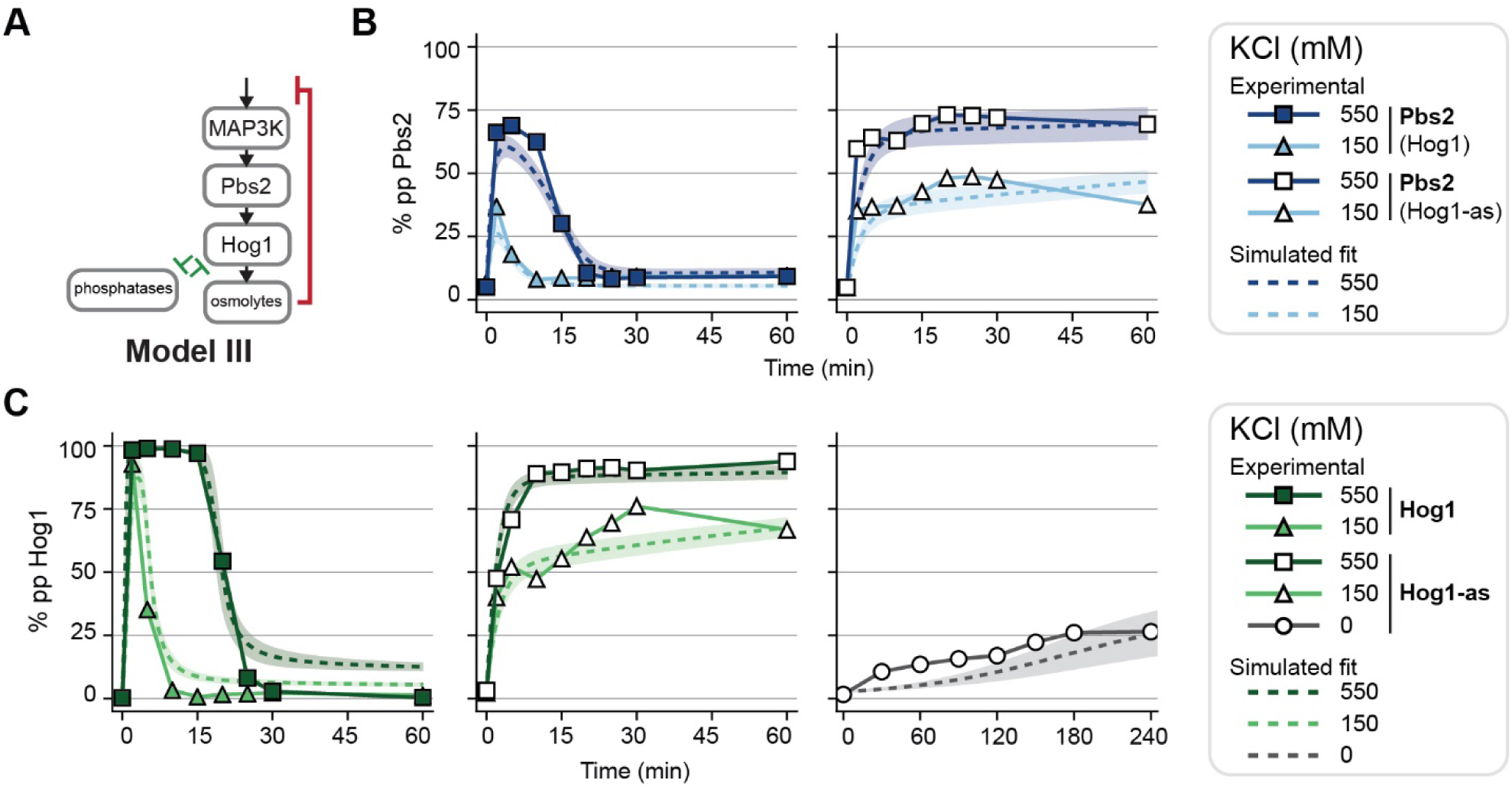
Model III with positive feedback acting through mutual inhibition captures experimental data. **(A)** Schematic of Model III with a delayed negative feedback and a positive feedback decreasing its phosphatases’ activity. **(B)** Model III simulated Pbs2 fits (dashed lines) overlaid with data (symbols). Left: Data and simulations for wildtype Hog1 in response to 550 mM and 150 mM KCl. Right: Data and simulations for Hog1-as. **(C)** Model IIa simulated Hog1 fits (dashed lines) overlaid with data (symbols). Left: Data and simulations for wildtype Hog1 in response to 550 mM and 150 mM KCl. Center: Data and simulations for Hog1-as. Right: Data and simulations for Hog1-as with no salt stimulus. All simulations are n = 1000 and shaded regions represent a SD of 1.

**Figure 7 – figure supplement 2:**
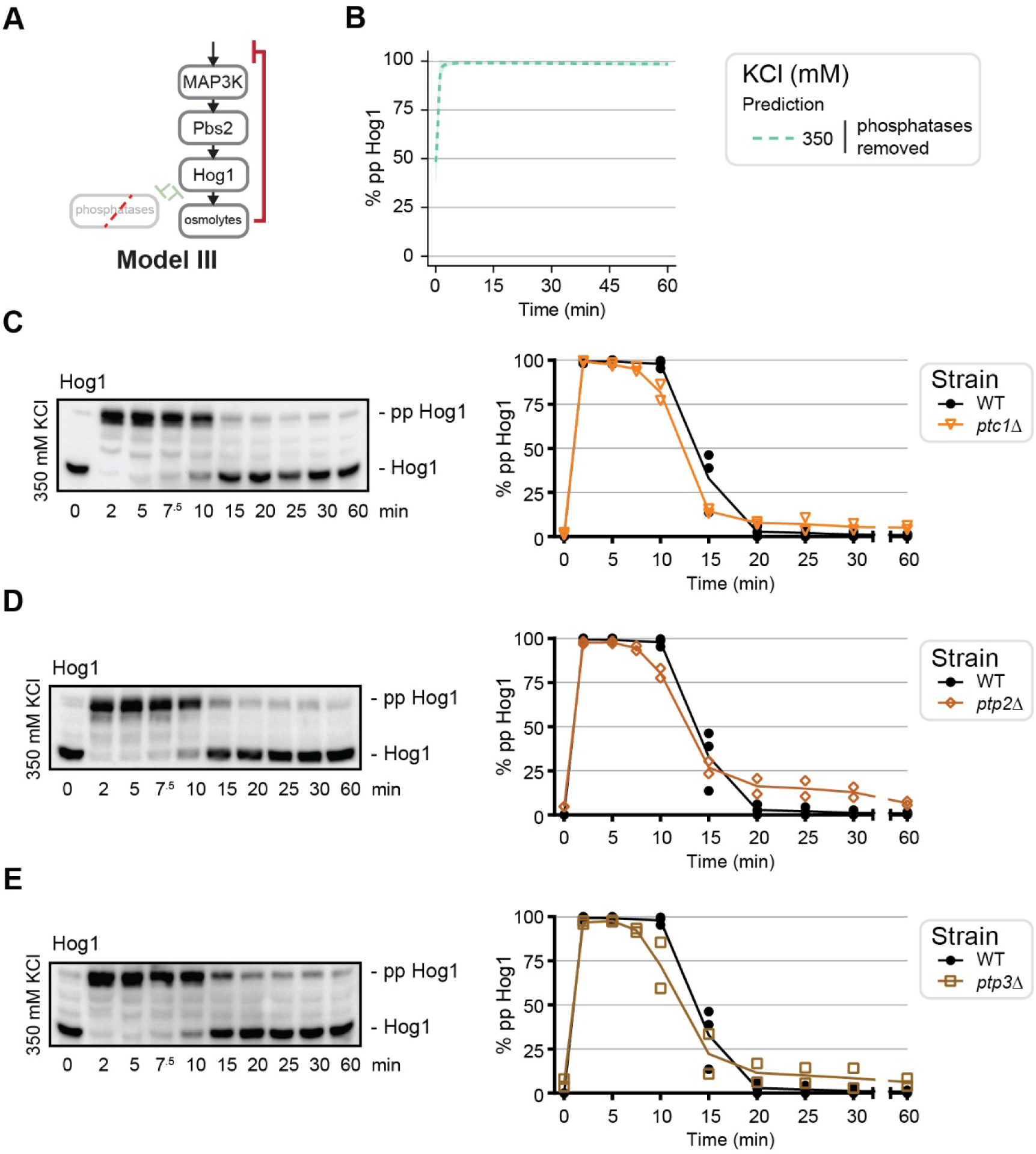
Single deletions of the primary Hog1 phosphatases slightly decrease duration of Hog1 activation. **(A)** Schematic of Model III with mutual inhibition acting as positive feedback, here depicted as the phosphatases removed. **(B)** Model III Hog1 prediction in response to 350 mM KCl if the phosphatases were removed. Simulations are n = 1000 and shaded regions represent a SD of 1. **(C)** Left: Hog1 behavior in response to 350 mM KCl in *ptc1*Δ background. Right: Quantification of blots. **(D)** Same as (A) for *ptp2*Δ background. **(E)** Same as (A) for *ptp3*Δ background.

